# Identification of novel HDAC11 inhibitors: *In silico* & *in vitro* studies

**DOI:** 10.64898/2026.08.24.746593

**Authors:** Mohon Paul, Devika S Kumar, Sujata Mishra, Arunasree M Kalle

**Affiliations:** Department of Animal Biology, School of Life Sciences, University of Hyderabad, Gachibowli, Hyderabad, TS, India 500046

**Keywords:** HDAC11, Selective inhibitors, Zinc chelation, Homology modeling, Molecular dynamics

## Abstract

Histone deacetylases (HDACs) are pivotal epigenetic regulators that modulate diverse cellular pathways by removing acetyl groups from lysine residues on both histone and non-histone proteins. Histone deacetylase 11 (HDAC11), the sole member of class IV HDACs, exhibits both deacetylation and fatty acid deacylation activities. Accumulating evidence implicates HDAC11 as a key epigenetic regulator of fundamental cellular processes, including metabolism, immune responses, and tissue development. Dysregulation of HDAC11 activity has been associated with inflammatory diseases, metabolic disorders, neurodegenerative conditions, and cancer, highlighting its potential as a therapeutic target. Although several HDAC11-specific inhibitors have been identified, none have progressed to clinical development. In this study, we aimed to discover HDAC11-selective inhibitors by integrating in silico and in vitro validation approaches. Homology modelling of the HDAC11 structure was conducted, followed by model validation, structure-based virtual screening, molecular dynamics (MD) simulations, and binding free energy calculations. We identified and validated three lead compounds and their intermediates using biochemical and cell-based assays. Fluorescence-based and HPLC-based enzymatic assays demonstrated potent inhibition of both the deacetylase and deacylase activities of HDAC11, with Inhibitor 6 and Inhibitor 3 exhibiting the strongest effects among the six compounds tested. Further, a decrease in lipid accumulation, reduced stability of the HDAC11 substrate SHMT2, as determined by immunoblot analysis and decreased cell viability, as assessed by MTT assay, confirmed HDAC11 inhibition in cellular models. The study shows that new HDAC11 inhibitors significantly reduce the viability of breast cancer cells and induce apoptosis; inhibitor 6, in particular, showed high potency, similar to the reference compound SIS-17. Flow cytometry showed that treated MDA-MB-231 cells exhibited cell-cycle arrest and increased apoptosis, a finding further confirmed by Annexin V/PI staining. Molecular analysis showed that BAX increased while BCL2 decreased, indicating that apoptotic pathways were activated in novel compound-treated MDA-MB-231 cells. The results suggest that inhibiting HDAC11 is an effective way to induce cancer cell death and provide a basis for further assessment of these compounds as potential treatments for breast cancer. Collectively, this study identifies novel zinc-chelating HDAC11 inhibitors containing a nitro-sp² group, providing promising candidates for further therapeutic development.

## Introduction

Gene expression is regulated by epigenetics, which occurs without altering the DNA sequence. This control involves processes such as DNA methylation, histone modifications, and chromatin remodelling [3–5]. Among these, histone acetylation and deacetylation are reversible modifications mediated by histone acetyltransferases (HATs) and histone deacetylases (HDACs), respectively, which govern transcriptional activation or repression [6,7]. The 18 isoforms of HDACs known in humans, play essential roles in maintaining cellular balance by regulating both histone and non-histone substrates [8–13] *via* lysine deacetylation.

HDAC11 is the most recently discovered and smallest member of HDAC family, possessing deacetylation and deacylation activities. HDAC11 has been increasingly implicated in metabolic disorders, particularly diabetes [12,13]. Mechanistic studies suggest that HDAC11 overexpression contributes to insulin resistance by disrupting insulin receptor signaling and promoting the release of inflammatory cytokines in adipose tissue [12,14]. It has been linked to regulating lipid metabolism and energy homeostasis in fat and liver diseases. Its regulation of fatty-acylation also affects lipid handling, leading to altered adipogenesis and ectopic fat accumulation [15,16]. Moreover, HDAC11 activity has been associated with impaired β-cell function and reduced glucose-stimulated insulin secretion, both distinct features of type 2 diabetes [17]. It regulates the interaction of interleukin-10 with metabolic enzymes such as SHMT2 [17].

Inhibition of HDAC11 has shown promise in restoring insulin sensitivity, improving glucose tolerance, and mitigating obesity-associated metabolic dysfunction, thereby positioning it as a novel therapeutic target [18]. In cancer, loss of HDAC11 expression suppresses tumour growth, inhibits immune evasion and establishment, and promotes cell death by apoptosis, making HDAC11 a potential epigenetic biomarker and drug target in multiple cancer types [18].

Several HDAC inhibitors (HDACi) are in clinical usage for various diseases including Vorinostat (SAHA) and Romidepsin (FK-228) for T-cell lymphoma, Belinostat for peripheral T-cell lymphoma, and Panobinostat for multiple myeloma [11,19]. Although effective, these broad-spectrum inhibitors (pan-HDACi) often cause adverse effects such as fatigue, gastrointestinal disturbances, thrombocytopenia, and neutropenia due to their lack of isoform specificity [7,20]. This highlights the importance of developing HDAC-selective inhibitors that minimize toxicity while preserving therapeutic benefits. Hydroxamic acid–based HDAC11 inhibitors (e.g., FT895, Elevenostat, and SIS17) and macrocyclic peptide inhibitors (e.g., TD034) have shown anticancer potential, but none of them are in clinic yet and require further optimization to develop HDAC11 isoform-selective molecules[19,21].

In this study, computational approaches such as structure-based drug design, pharmacophore modelling, molecular docking, and molecular dynamics (MD) simulations were employed to discover novel HDAC11 inhibitors and the selected candidate compounds were further assessed using *in vitro* cellular assays.

## Results

### Homology Modelling and Active Site Characterization

The 3D model of HDAC11 was constructed with zinc coordination and evaluated for quality against the SAVES6.0 server (**Fig. 1A**). The model achieved an Errat quality score of 96.1. The Ramachandran analysis showed that 89.8% of the residues were in the most favored region. Only 0.4% of residues were in the disallowed region, potentially indicating that the structural quality of the model is adequate (**Fig. 1B**). HDAC11’s active site was compared with those of HDAC4, HDAC6, and HDAC8 to identify key differences and similarities. The analysis revealed several conserved residues crucial for inhibitor binding: Y304: Hydrogen bond donor to ligand hydroxamate, H142–H143: Donor–acceptor pair. F152, Y209, H183: Hydrophobic stacking interactions, Asp174, Asp254, His176: Zinc coordination. PLIP and LigPlus confirmed zinc coordination within the active site, ensuring proper positioning for catalytic activity. Based on these interactions and using PharmaGist, we subsequently designed five pharmacophore models (**Table 1**), guided by ligands including known HDAC11 inhibitors FT895, SIS17, TSA, and Elevenostat. The models varied in the number and arrangement of hydrogen bond donors and acceptors to capture diverse binding modes (**Fig. 1C**).

**Figure 1.**
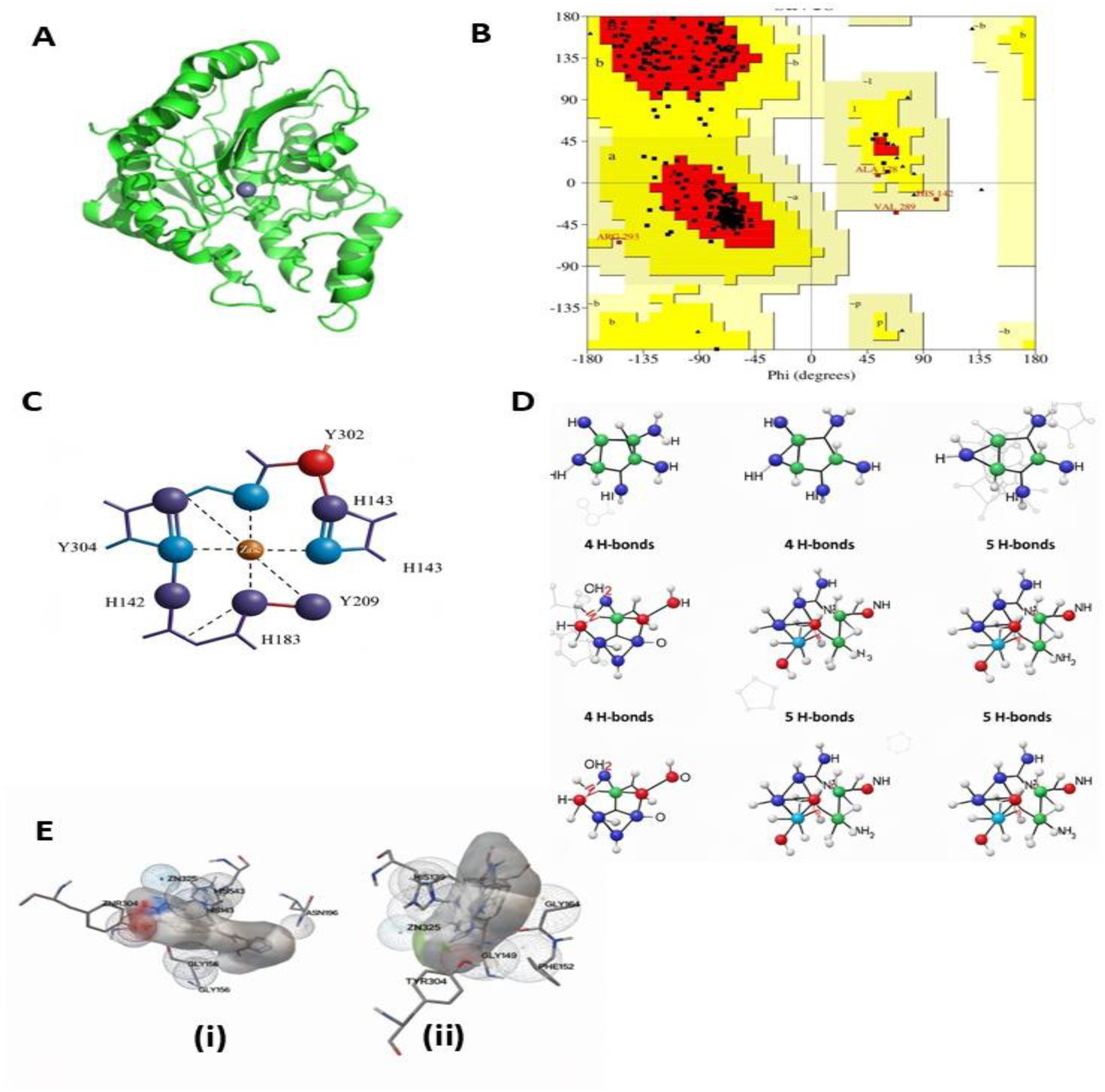
Homology Modelling of HDAC11. (A) Structure of HDAC11 (B) Ramachandran Plot of HDAC11 (C) **Key Active Site Residues**: **Y304**: Hydrogen bond donor to ligand hydroxamate, **H142–H143**: Donor–acceptor pair, **F152, Y209, H183**: Hydrophobic stacking interactions, **Asp174, Asp254, His176**: Zinc coordination. (D) Pharmacophore models were generated based on known HDAC inhibitors: FT895, SIS17, Elevenostat, and TSA. (E) i. Interaction of HDAC11-Elevenostat complex, ii. Interaction of HDAC11-FT895 complex

**Table 1.**
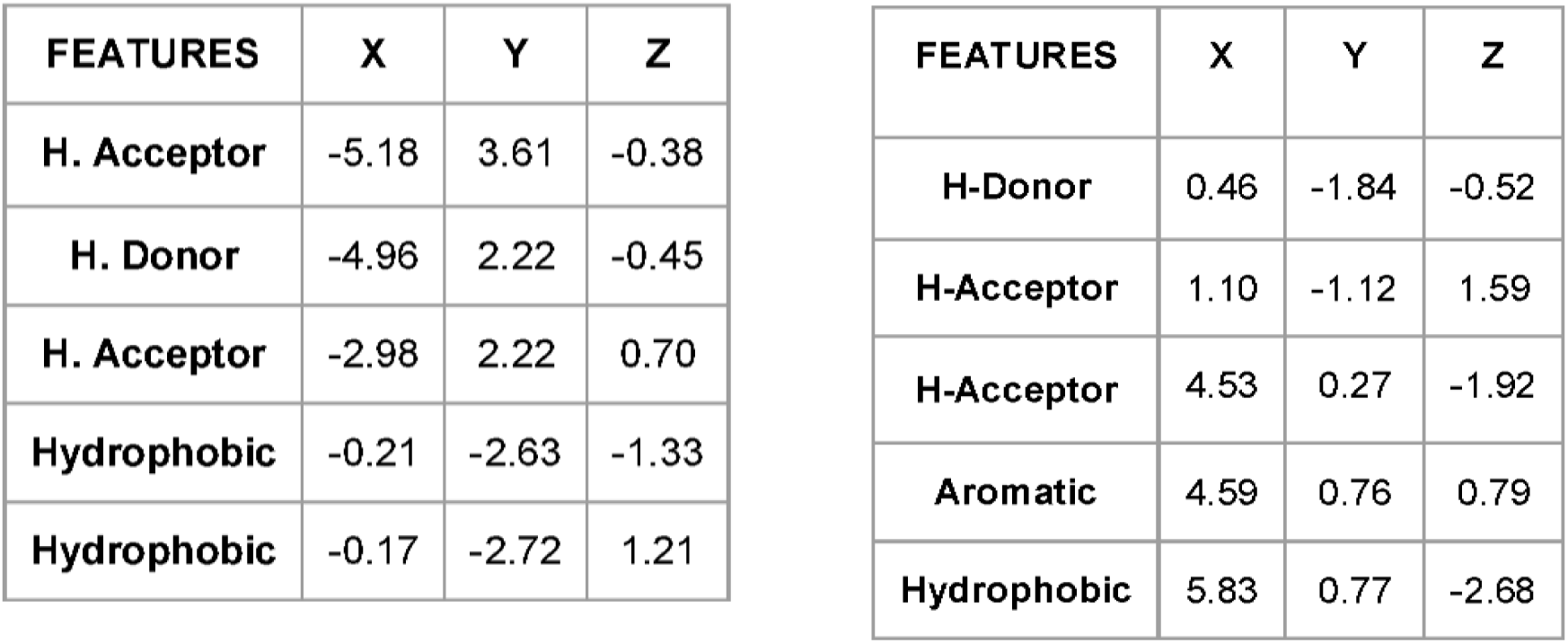
MD simulation analysis of ZINC30713663-HDAC11 complex.

### Docking, Molecular Dynamics Simulations, and Analysis

The five pharmacophore models were used to screen the ZINC database, a large collection of commercially available compounds. ZincPharmer was used for the virtual screening process and the compounds matching the pharmacophore features were selected as potential hits for further docking. Molecular docking was performed using PyRx, a virtual screening tool. The AutoDock4Zn protocol was used to account for the zinc ion in the active site and a grid box centered on the zinc ion (25 Å³) was used to define the search space for ligand binding. Approximately 76,000 ligands were docked into the HDAC11 active site and using LigGrep compounds that interacted with Zn and Y304 were selected as lead molecules and were further refined for drug-like properties using DruLiTo. A total of 30 lead molecules were then redocked using Smina with the Lin_F9 scoring function to improve the accuracy of binding affinity predictions. Interactions between the top five ligands and HDAC11 were analyzed using BINANA, PLIP, and PoseView to identify key binding modes and interactions. (**Supplementary File 1 Fig. 1**). Furthermore, PLIP analysis confirmed zinc chelation *via* the nitro-sp2 group of the ligands, which is crucial for HDAC11 inhibition.

Known HDAC11 inhibitor, FT895 was employed for performing MD simulations and confirm the predicted active-site residues and previous structural findings. The simulations provided valuable insights into ligand binding and structural stability. The MD simulation results were processed and analyzed using the CPPTRAJ module from AmberTools. Among the tested compounds, ZINC13619108, ZINC15938334, and ZINC24080467 exhibited relatively stable performance in terms of RMSD, RMSF, hydrogen bonding, and radius of gyration. In contrast, ZINC12560584 showed significant fluctuations in these parameters, while ZINC30713663 demonstrated notably high RMSD, RMSF, and radius of gyration values. Notably, ZINC13619108, ZINC24080467, and ZINC30713663 produced positive MMPBSA results, which are unfavorable. The MMGBSA value of the ZINC15938334-HDAC11 complex was closer to and slightly lower than that of the FT895-HDAC11 reference complex. On the other hand, ZINC12560584, ZINC24080467, and ZINC13619108 showed highly negative MMGBSA values, whereas ZINC30713663 had negative results, though not within the threshold for a potent inhibitor (**Table 2**).

**Table 2.**
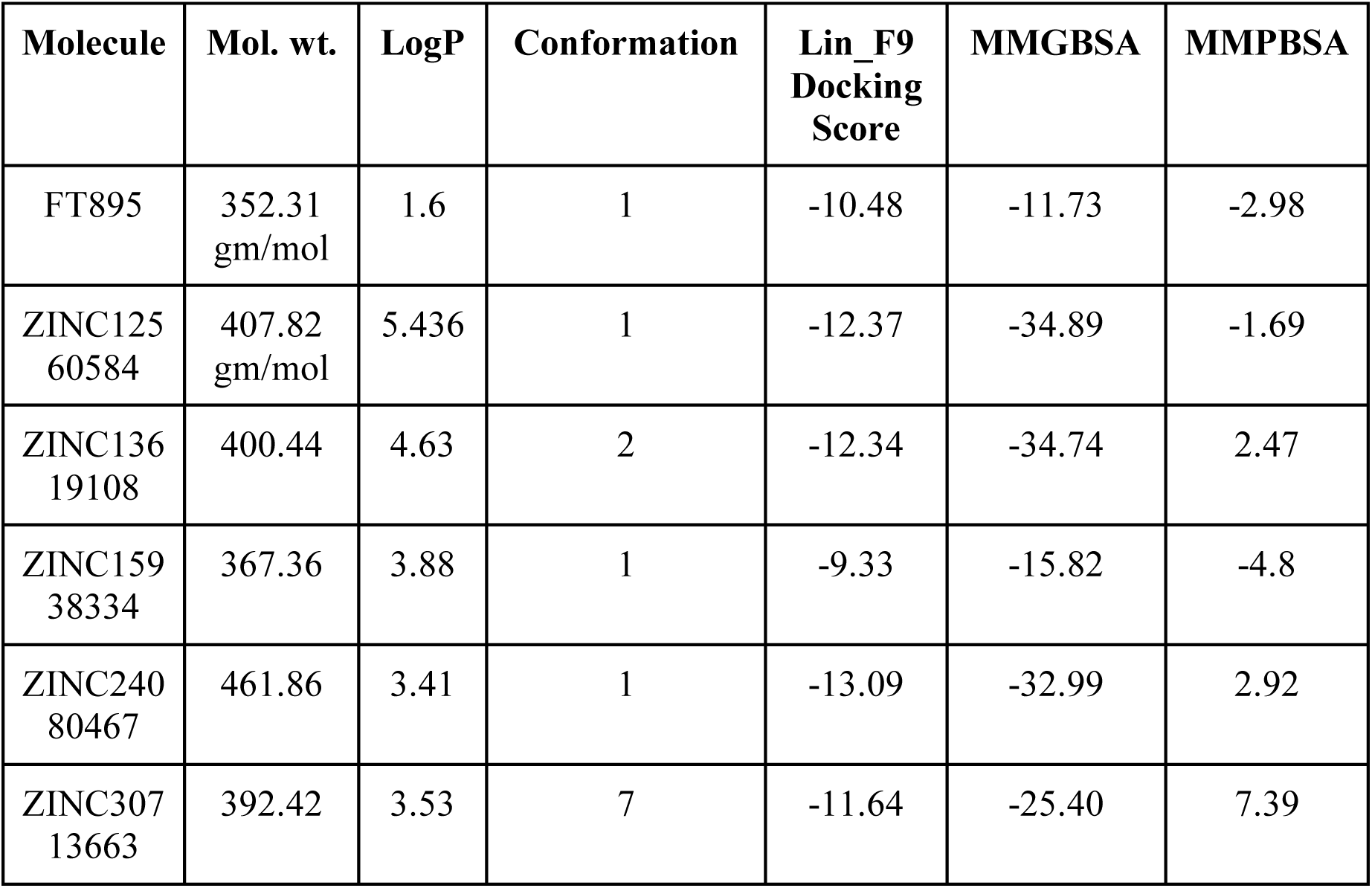
Binding Free Energy Analysis.

ZINC13619108, ZINC15938334, and ZINC24080467 showed the most stable interactions with the HDAC11 active site and therefore these three compounds along with their major intermediates **(FIG 3)** were synthesized (**Supplementary File 1 Fig. 2)** and characterized by NMR and LC-MS (**Supplementary File 2**).

**Figure 2.**
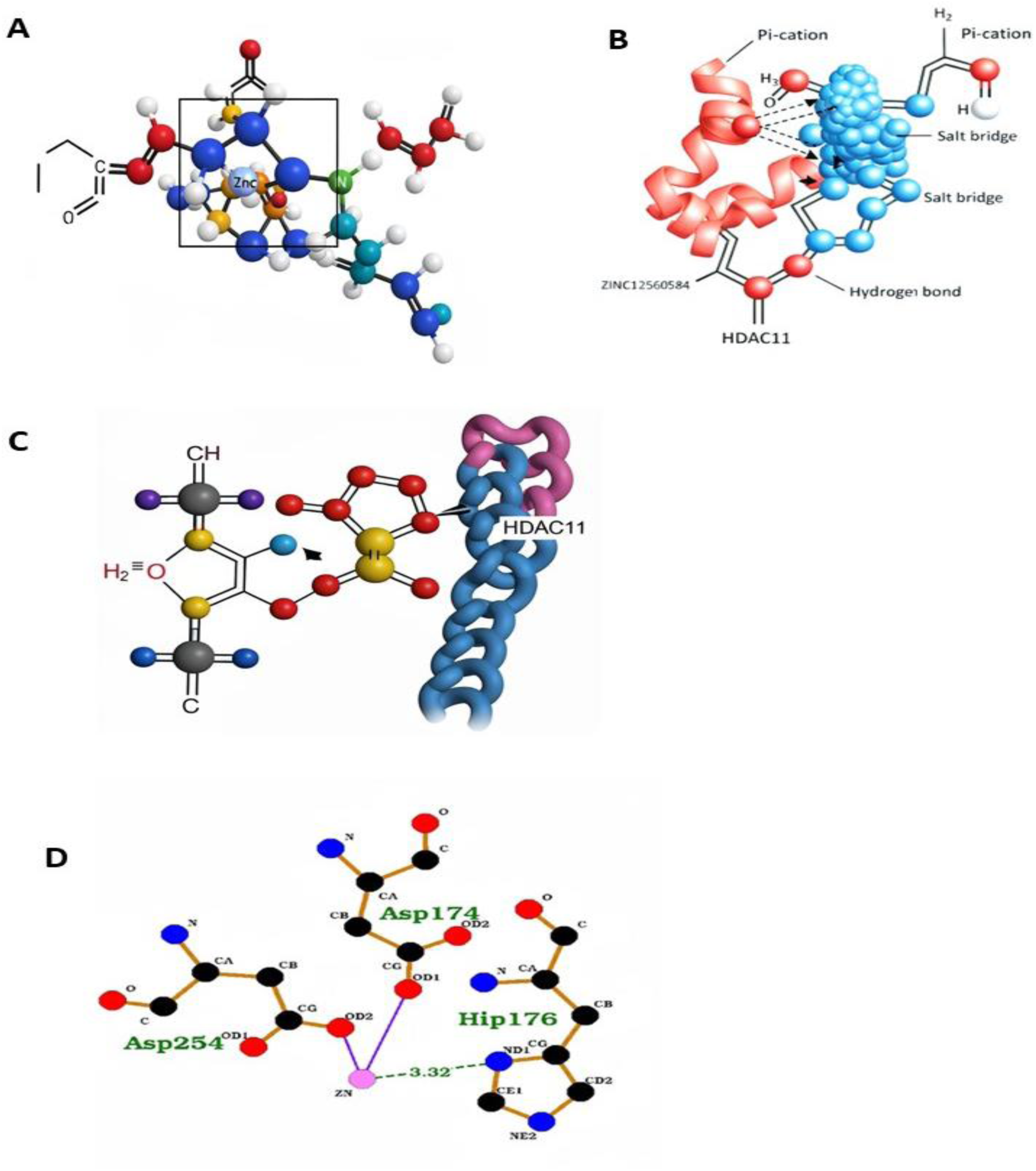
Molecular Docking. (A) HDAC11 Docking with PyRx (B) Model Validation with Known Inhibitors (C) Zinc chelation via the nitro-sp2 group of the ligands confirmed by PLIP analysis (D) The interactions with Zinc and the standalone HDAC11 molecule

**Figure 3.**
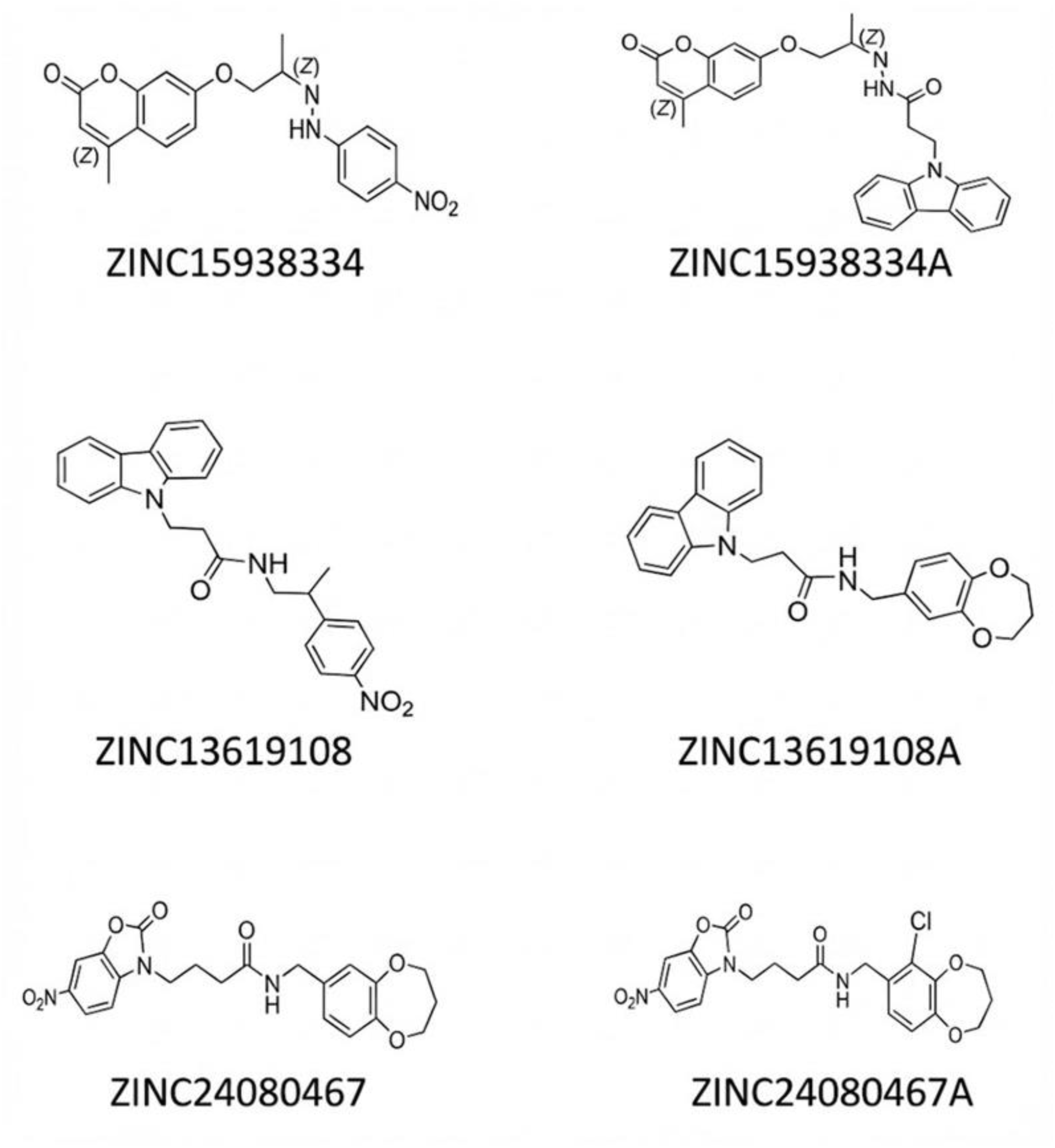
Structure of selected molecules for MD simulation and their reaction precursors.

**Figure 4.**
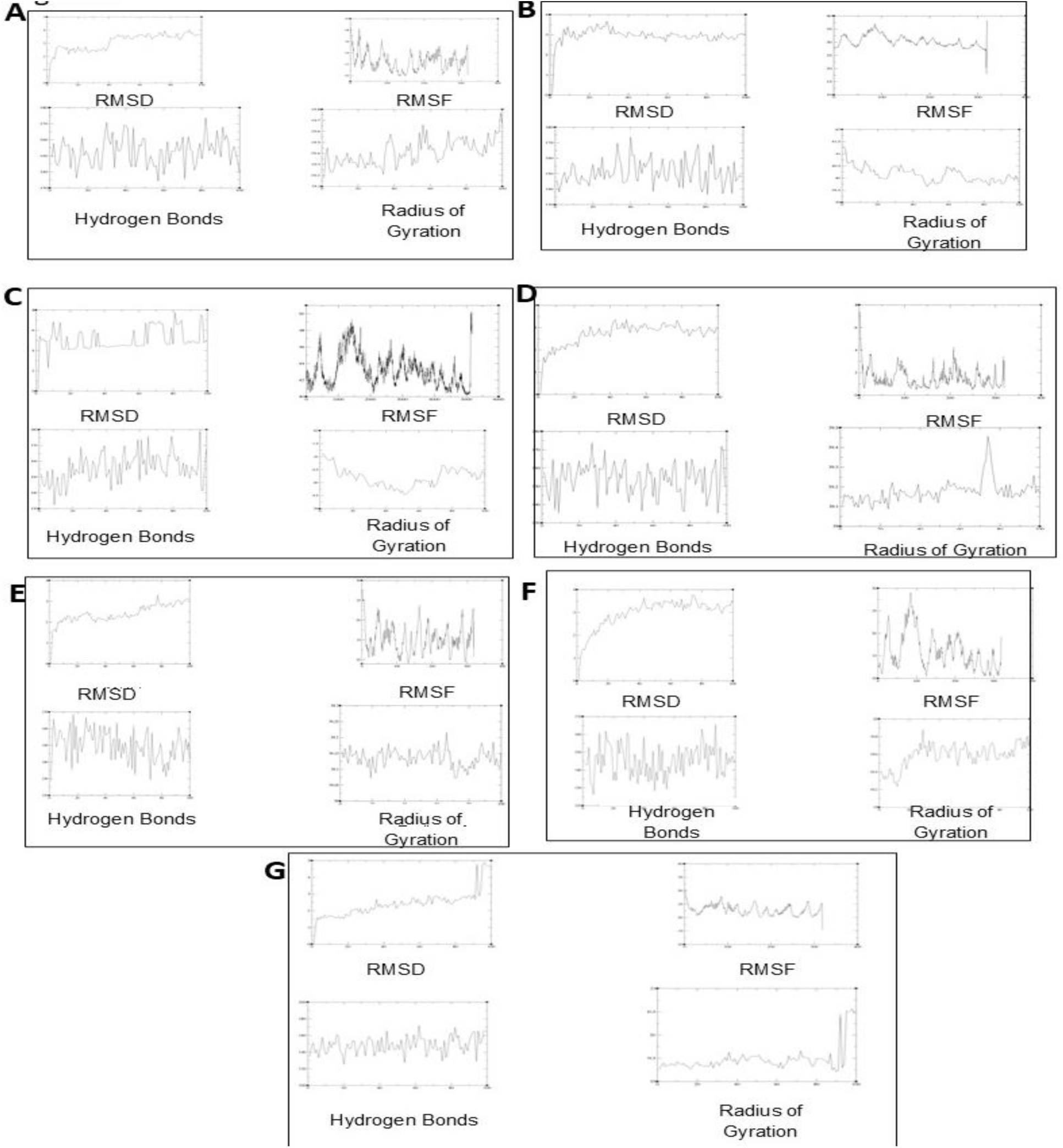
The RMSD, RMSF, Hydrogen bond, and Radius of Gyration analysis of the standalone HDAC11 protein. (A) Standalone HDAC11 protein (B) FT895-HDAC11 Complex (C) ZINC12560584-HDAC11 complex (D) ZINC13619108-HDAC11 complex (E) ZINC15938334-HDAC11 complex (F) ZINC24080467-HDAC11 complex (G) ZINC30713663-HDAC11 complex

### The selected compounds inhibited the deacetylase and deacylase activity of HDAC11

The deacetylase inhibitory activity was of the 6 compounds was assessed using immunoprecipitated HDAC11 and crude lysate as enzyme source and acetylated lysine substrate. Comparing the crude lysate enzyme activity with the HDAC11 immunoprecipitated activity, Inhibitor 6, Inhibitor 3 followed by Inhibitor 4 showed significant HDAC11-specific deacetylation inhibitory activity comparable to the known HDAC11 inhibitor, SIS-17 (**Fig. 5A**). Inhibitor 1 & 2 showed least HDAC11-specific inhibitory activity comparable to pan-HDAC inhibitor SAHA suggesting non-specific HDAC inhibition of those two compounds.

**Figure 5.**
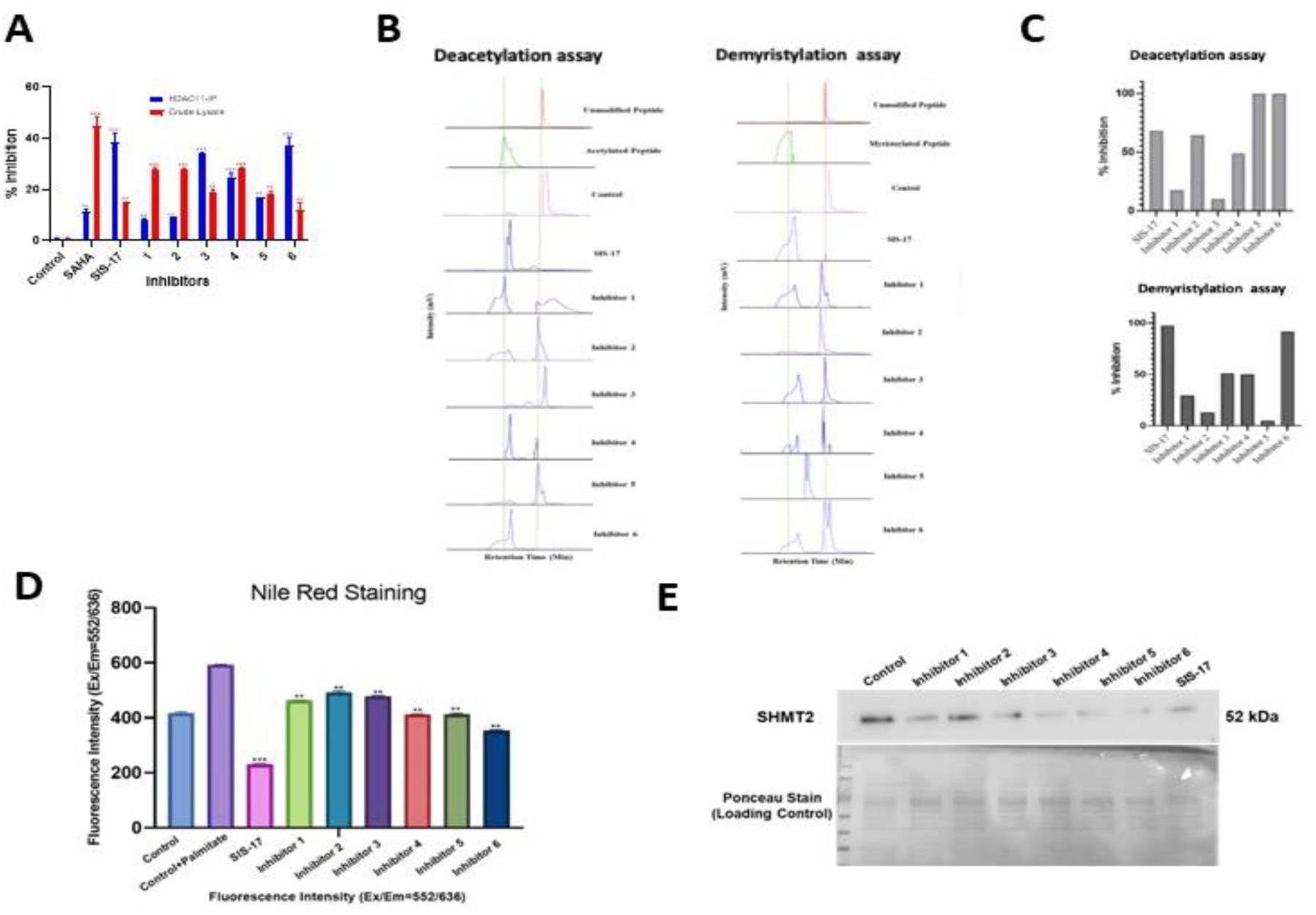
In-vitro Validation of the selected molecules. (A) HDAC11 deacetylase activity assessed using immunoprecipitated HDAC11 and crude lysate (B) HPLC-based enzymatic peptide assay to assess both deacetylase and deacylase activity of HDAC11 (C) Quantification of the HPLC-based enzymatic peptide assay (D) The lipid accumulation was quantified by Nile Red Assay in HepG2 cells (E) Immunoblot analysis of SHMT2 in MCF7 cells treated with or without the selected compounds (F) Cell viability assay using MTT in MCF-7 breast cancer cells treated with the inhibitors at different concentrations1(nM to 100µM) for 24 hours. *p< 0.05, **p < 0.01, ***p<0.001 and ****p<0.0001 compared to the control.

To further evaluate the deacetylation and demyristylation activity of HDAC11, HPLC-based enzymatic peptide assay was performed using acetylated and myristylated lysine peptides and immunoprecipitated HDAC11 protein. The assay showed that the inhibitors affected both the demyristylation and deacetylation activity of the enzyme (**Fig. 5B**). Inhibitor 6 showed complete inhibition of HDAC11 activity and is at par with the known inhibitor, SIS-17. Inhibitors 3 and 4 showed more than 50% inhibition of demyristylation activity of HDAC11, however, inhibitor showed 50% inhibition of deacetylation activity suggesting inhibitor 3 is a more potent HDAC11 demyristylation inhibitor (**Fig. 5C**).

### HDAC11 inhibition resulted in decreased lipid accumulation and SHMT2 stability

HepG2 liver cancer cell line was subjected to treatment first with BSA-Palmitate to mimic the lipid accumulation condition and subsequently treated with the 6 novel inhibitors along with SIS-17 for 24 hours. The lipid accumulation was quantified by Nile Red Assay. Inhibitor 6 showed reduced lipid accumulation suggesting inhibition of HDAC11 (**Fig. 5D**). To further assess the HDAC11 inhibitory activity by the compounds, the protein levels of HDAC11 substrate, SHMT2, were analyzed by immunoblot analysis in MCF7 cells treated with or without the compounds. The SHMT2 levels were reduced upon inhibitor treatment suggesting HDAC11 inhibition destabilizes the myristylated SHMT2 marking for proteasomal degradation (**Fig. 5E**). Inhibitor 6 showed increased HDAC11 inhibition as evident from the decreased SHMT2 levels (**Supplementary File 1 Fig. 3**).

### Inhibition of HDAC11 has potential anti-cancer effects

The MCF-7 breast cancer cells were seeded and treated with the inhibitors at different concentrations (1nM to 100µM) for 24 hours (**Fig. 6A**). The cell viability assay using MTT showed a characteristic decrease in the percent cell viability of the cells upon treatment with increasing concentration of all the inhibitors. Inhibitor 6 showed potent anti-cancer activity on MCF7 cells at par with the SIS-17, with lower IC_50_ concentration.

**Figure 6.**
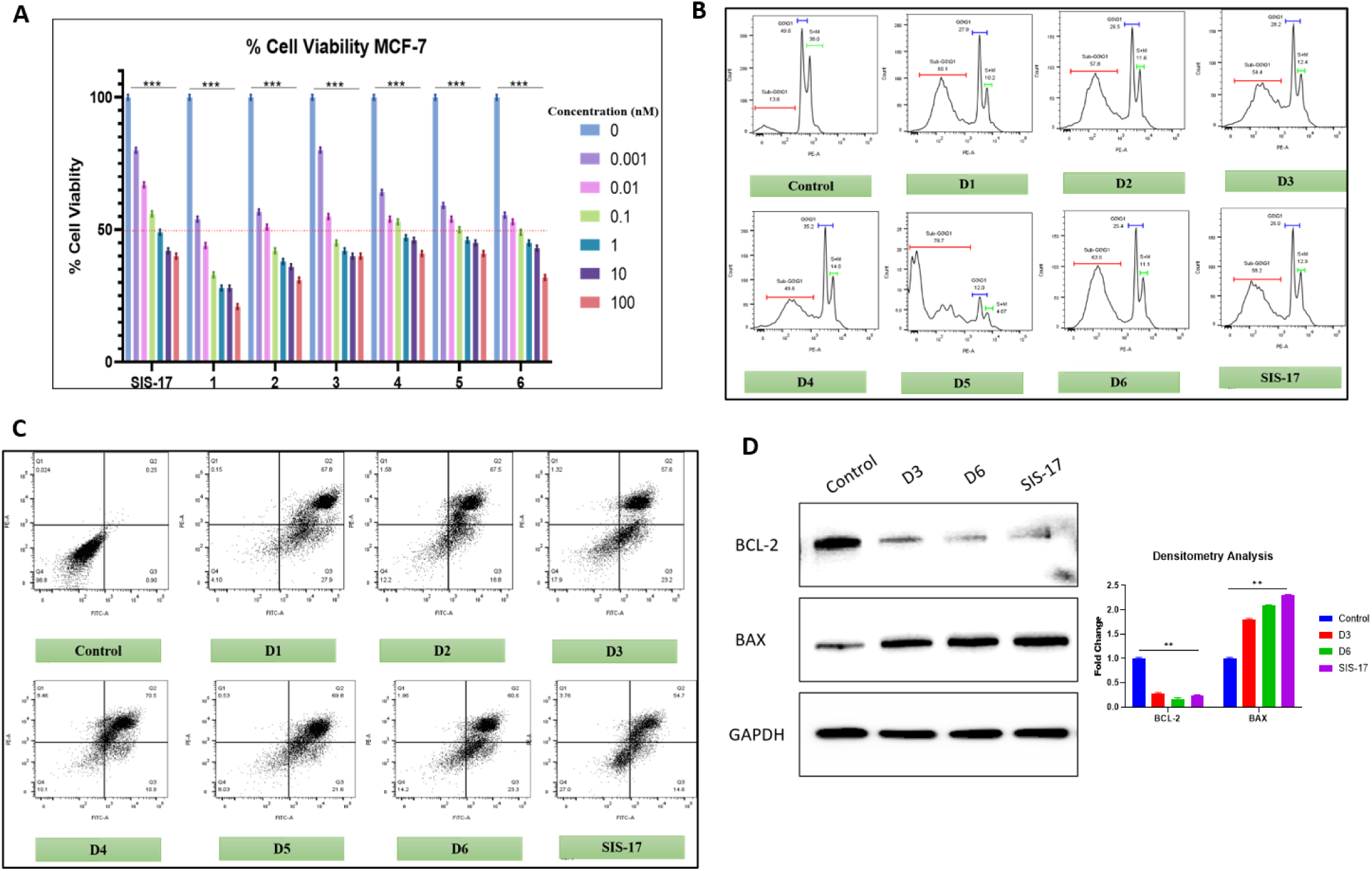
HDAC11 Inhibitors Induce Cell Cycle Arrest and Apoptosis in Breast Cancer Cells. (A) Dose-dependent cytotoxicity of HDAC11 inhibitors in MCF7 cells. All inhibitors reduced viability, with Inhibitor 6 showing the greatest potency (lowest IC₅₀), similar to SIS-17. (B) Cell cycle analysis of MDA-MB-231 cells after HDAC11 inhibitor treatment. Treated cells showed increased Sub G0-G1 population and G2/M arrest compared to control. (C) Apoptosis induction in MDA-MB-231 cells measured by Annexin V/PI staining. Inhibitor 6 strongly increased early and late apoptotic populations. (D) Western blot analysis of BAX and BCL2 in MDA-MB-231 cells. Effective inhibitors increased BAX, decreased BCL2, and raised the Bax/Bcl-2 ratio.

To understand how growth was inhibited, flow cytometry was used to analyze the cell cycle in MDA-MB-231 cells treated with new compounds (Drugs 1–6), the positive control SIS-17, and an untreated control (**Fig. 6B**). In the control group, most cells were in the G0/G1 phase, showing a normal cell cycle. When treated with SIS-17 and the new HDAC11 inhibitors, there was a clear shift in cell cycle phases. More cells appeared in the Sub G0-G1 phase than in the control, suggesting a G2/M cell-cycle arrest.

To measure apoptosis, the treated cells were stained with FITC Annexin V and PI. The untreated control cultures remained highly viable, with over 90% of cells in the double-negative quadrant (**Fig. 6C**). Among the compounds, Inhibitor 6 showed strong pro-apoptotic effects, with many cells moving into both the early and late apoptotic quadrants. This shows that the cytotoxic effect might be due to apoptosis via regulated pathways.

Further, immunoblot analysis was performed to measure the expression levels of apoptotic proteins (**Fig. 6D**). Specifically, the pro-apoptotic protein BAX and the anti-apoptotic protein BCL2 were probed. In untreated control cells, BAX and BCL2 were at baseline levels. When treated with Inhibitor 3, Inhibitor 6, and SIS-17, BAX protein levels increased significantly, while BCL2 levels decreased. Statistical analysis using quantitative densitometry showed a significant increase in the overall Bax/Bcl-2 ratio in groups treated with effective compounds. These results confirm that the new compounds cause cell death in MDA-MB-231 cells.

## Discussion

HDACs are known to epigenetically regulate several cellular pathways by lysine deacetylation of histone and non-histone proteins [14]. Aberrant expression of HDACs cause several diseases such as inflammation, neurodegenerative diseases and cancer [7]. Pan-HDACi such as Vorinostat, Panobinostat, Belinostat etc. have proven efficacy as anti-cancer activities [19] however their usage is limited by severe adverse effects due to non-specific HDAC inhibition. Class-selective, isoform-specific HDAC inhibitors have greater therapeutic value and are the current research focus.

Histone deacetylase 11 (HDAC11), the sole member of class IV HDACs, has recently been identified as a significant regulator of lipid metabolism, immune modulation, and tumor progression [13–15]. HDAC11 is thought to have evolved recently and has shares lower sequence and structural similarity with other HDACs. Gene knockout studies in mice have clearly demonstrated that HDAC11 deficiency improves metabolic diseases such as diabetes and high-fat induced obesity [12,13].Although few potent HDAC11 inhibitors have been developed in with preclinical studies, such as FT895 and SIS17, there are no clinically approved HDAC11-specific inhibitors; thus, this study aimed to identify potent HDAC11 inhibitors using *in silico* approaches, synthesize and characterize the hit molecules and biologically assess their potency and efficacy using *in vitro* assays.

Crystal structure of HDAC11 is unavailable and therefore, we modelled HDAC11 structure using homology modelling where HDAC6 catalytic domain 2 (CD2) was used as template due to its higher homology with HDAC11. Standard docking programs such as AlphaFold and SWISS-MODEL which predicted and modelled HDAC11 structure, do not include metal ion such as Zn and fail to provide accurate interactions and binding free energies. [21,23] A zinc ion was manually added to the active site by structural alignment with other zinc-dependent HDACs. Active site residues analysis provided meaningful structural information about the catalytic pocket of HDAC11. The homology model contained conserved hydrophobic domains and zinc-binding domains, typical in all HDAC enzymes, validating the structure-based screenings.

Virtual screening and molecular dynamics (MD) simulations identified a group of compounds with high binding affinities and stable interactions between ligands and catalytic residues within the active site. Notably, the stability of the binding could be attributed to coordination with the zinc ion or formation of hydrogen bonds. In addition, these observations indicate the structure’s reliability for identification using *in silico* predictions, guiding the selection of compounds for biochemical screening. We finally identified three lead molecules and their intermediates based on their favorable docking score, MMGBSA, MMPBSA, and stability for further in-cell validation.

Biochemical assays demonstrated that the selected compounds inhibited HDAC11 deacetylase activity, in accordance with the docking inhibition profile. Additionally, the HPLC-based enzyme assays showed that the compounds inhibited HDAC11 fatty deacylase activity, as indicated by a shift in the peaks for both acetylated and myristoylated peptides when treated with the inhibitors, particularly Inhibitor 6 and Inhibitor 3. This dual inhibition is especially relevant as HDAC11-mediated deacylation has been implicated in regulating metabolic pathways and protein stability [24,25] and its deacetylation in immune modulation [24] and tumor progression [26]. The Nile Red assay indicated that inhibition of HDAC11 resulted in decreased lipid accumulation, and the immunoblotting showed a decreased stability of SHMT2, a known substrate of HDAC11 and the results are in agreement with earlier studies [17]. The novel compounds exhibit potential anti-cancer effects, as observed by the dose-dependent reduction in cell viability of MCF-7 breast cancer cells, and an increased apoptosis in MDA-MB-231 cells. In addition, western blot analysis confirmed increased BAX/BCL2 ratio in control vs treated MDA-MB-231 cells, supporting the apoptotic signaling induced by the novel HDAC11 inhibitors. Overall, this *in silico* and *in vitro* integration study identifies HDAC11 inhibitors with Zinc chelation using novel Nitro-sp2 group. Future studies will focus on lead optimization and testing the efficacy in disease-associated models.

## Methodology

### In Silico Studies

#### Model Construction and Validation

Since the HDAC11 protein structure was not available, the human HDAC11 cDNA sequence (UniProt ID: Q96DBZ) was used to create a homology model by the Swiss-Model server, taking the catalytic domain 2 (CD2) of HDAC6 (PDB ID: 5EDU) as a structural template. The first model was refined and optimized using ModRefiner and DeepRefiner, and the zinc ion from the template was incorporated into the structure of HDAC11. The model was then minimized for energy using the YASARA server, and the three-dimensional model was generated in a final PDB format. The quality, structure and model were verified through the SAVES6.0 server [27–31].

#### Active Site Mapping

The active-site residues of HDAC4, HDAC6, and HDAC8 were identified from ligand-bound crystal structures present in the Protein Data Bank using LigPlus along with some references from the literature. Structural alignments with HDAC11 and interaction profiles from the PLIP server were further used to identify conserved residues important for HDAC11 activity.

#### Pharmacophore Modelling and Virtual Screening

The three established HDAC11 inhibitors, FT895, SIS17 and Elevenostat, and pan-HDAC inhibitor, TSA, were utilized to generate five pharmacophore models using PharmaGist to cover different arrangements of hydrogen bond acceptors and donors, which were then used for virtual screening within ZincPharmer for ligands that map onto HDAC11.

#### Molecular Docking

Docking studies of the HDAC11 model were conducted with the AutoDock tools for the preparation of the receptor and PyRx for the simulations. A docking grid was centered on the zinc ion with the dimensions of 25 Å³ and exhaustiveness of 32. The molecules were further filtered through LigGrep on the interactions with the zinc ion and the active-site residue Y304. The lead molecules were evaluated for drug-likeness *via* DruLiTo with Lipinski, Ghose, and Q-Filtering. Selected compounds were further docked using Smina using Lin_F9 as the scoring function for the molecules. Interrogated profiling, hydrogen bonding, hydrophobic effects, salt bridges and π-stacking, was done with BINANA, PLIP, and PoseView. Five lead molecules were filtered down for further analysis by molecular dynamics.

#### Molecular Dynamics Simulations

AmberTools’ LEaP module with the SLEF force field was used to characterize the protein-ligand complexes. The system was minimized and equilibrated using PMEMD, and for the production runs, 100 ns of simulation was performed using PMEMD.cuda under periodic boundary conditions. Structural dynamics were analyzed using AmberTools’ CPPTRAJ to include RMSD, RMSF, radius of gyration, and hydrogen bonds. Selected molecules and their intermediates were synthesized from PozeSCAF Discovery Solutions Pvt. Ltd., India.

### In-cell Assays

#### Biochemical HDAC11 Enzyme Activity Assay (Kit-based)

To assess deacetylase inhibitory activity of the compounds, HDAC11 protein was immunoprecipitated from MCF7 cells as described previously [32]. The 6 novel inhibitors (10µM each), SIS-17 (10µM), along with the control (only protein), were added to HDAC11 and tested for enzyme activity using FLUOR DE LYS® substrate (Cat# BML-KI104-0050, ENZO Lifesciences, USA) as per manufacturer protocol.

#### HPLC-based Peptide Assay

The immunoprecipitated HDAC11 protein was used for performing an HPLC-based peptide assay [33]. The sequences of the peptide used were – KQTARKSTGGWW (Unmodified Peptide), KQTARK_(Ac)_STGGWW (Acetylated Peptide), and KQTARK_(Myr)_STGGWW (Myristoylated peptide). The HPLC was performed using Luna C18 HPLC Column (phenomenex®) using Buffer A HPLC-grade water with 0.1% trifluoroacetic acid and Buffer B HPLC-grade acetonitrile with 0.1% trifluoroacetic acid. The buffers were filtered using 0.22µm filter. The reaction buffer for the assay consisted of 20mM Tris-HCl (pH 8.0), 1mM DTT and 1mM NAD+. The assay was quenched using 20mM HCl and 320mM Acetic acid in Methanol. A standard curve for each peptide was generated with different peptide concentrations (1µM, 2 µM, 3 µM, 4 µM, 5 µM). The reaction setup of the peptide assay consisted of immunoprecipitated HDAC11 protein and peptide prepared in the reaction buffer for control, and presence of respective inhibitors in the test samples.

#### Mammalian Cell Culture

The human liver cancer cell line, HepG2 and breast cancer cell line, MCF7 were procured from the National Centre for Cell Science (NCCS, Pune, India). They were cultured in high-glucose Dulbecco’s modified Eagle’s medium (DMEM; GIBCO) supplemented with 10% fetal bovine serum (FBS; GIBCO) in the presence of 1% penicillin-streptomycin (GIBCO) at 37°C with 5% CO_2_ saturation.

#### Nile Red Staining

For lipid accumulation studies, HepG2 cells were cultured and seeded in 6-well plates at a density of 3 x 10^5^ cells/well. The cells were permitted to adhere overnight. Following this, cells were treated with 0.25 mM BSA-palmitate with or without the inhibitors mentioned above for 24 hr under the standard culture conditions. After incubation, cells were harvested and washed two times in 1X phosphate buffer solution (PBS). Cells were then stained for 10 min in the dark with 500 nM Nile Red (HIMEDIA). Excess dye was washed away using 1X PBS, and fluorescence was measured for determining intracellular lipid accumulation at excitation/emission wavelengths of 552 nm/636 nm using a fluorescence spectrophotometer.

#### Immunoblotting

The MCF-7 cells were subjected to a treatment with the six novel inhibitors, along with SIS-17 and a control (no treatment) for 24 hours. Proteins were isolated from the treated cells using RIPA (10 mM Tris-HCl (pH 7.4), 150 mM NaCl, 1% Triton X-100 or NP-40, 0.5% sodium deoxycholate, 0.1% SDS, and 1 mM EDTA) lysis buffer. 60 μg of total proteins were separated on 12% SDS-PAGE, immunoblotted and probed with anti-SHMT2 antibody (Cloud-clone Corp), anti-GAPDH antibody, anti-BAX antibody and anti-BCL2 antibody (SantaCruz). The protein bands were developed using chemiluminescence and visualized using the imaging system.

#### MTT Assay

MCF-7 cells were seeded in a 96-well plate at a seeding density of 5000 cells per well. They were subsequently treated with different concentrations (1nM, 10nM, 100nM, 1µM, 10 µM, 100 µM) of the inhibitors along with SIS-17 for 24 hours. The MTT assay was performed using 3-(4,5-Dimethylthiazol-2-yl)-2,5 Diphenyltetrazolium Bromide (MTT, HiMedia) as described earlier [32] and the absorbance values were recorded at 595 nm.

### Flow Cytometry Analysis of Cell Cycle and Apoptosis

MDA-MB-231 breast cancer cells were cultured in 6-well plates until they reached 70 to 80 percent confluence. The cells were then treated for 24 hours with either the vehicle, one of the new compounds (Drugs 1 to 6), or the positive control SIS-17. After treatment, both floating and attached cells were collected for analysis. To assess the cell cycle, DNA content was measured using Propidium Iodide (PI) staining. The collected cells were fixed overnight in cold 70% ethanol at 4°C, washed with PBS, and then incubated with PI solution containing RNase A. The samples were analyzed using a BD flow cytometer, and the distribution of cells in sub-G1, G0/G1, S, and G2/M phases was determined using established gating protocols [34]. To measure apoptosis, the FITC Annexin V Apoptosis Detection Kit (BD Biosciences) was used following the manufacturer’s instructions. After washing with cold PBS, the cells were resuspended in binding buffer and stained with FITC Annexin V and PI for 15 minutes at room temperature in the dark. The samples were then analyzed by flow cytometry. Cell populations were classified as viable (Annexin V−/PI−), early apoptotic (Annexin V+/PI−), late apoptotic (Annexin V+/PI+), or necrotic (Annexin V−/PI+).

### Statistical Tests

GraphPad Prism v8.4.3 software was used for the statistical analysis, which included a two-way ANOVA (two-way analysis of variance), and statistical significance (p-value) was established for * ≤0.05, ** ≤0.01, *** ≤0.001, **** ≤0.0001, and ns = non-significance ˃0.05.

## Author Contributions

MP performed the *in silico* work and drafted the manuscript. DSK performed the HPLC assay, immunoblotting, and flow cytometry; SM performed the biochemical assays. AMK conceptualized the study, arranged funding, analyzed the results, and reviewed the manuscript. All authors read and approved the final manuscript.

## Data Availability Statement

Data generated shall be made available upon request.

