## Supplementary data for "Identification of novel HDAC11 inhibitors: *In silico* & *in vitro* studies"

ZINC 13619108
1H CDCl3
16-09-2024

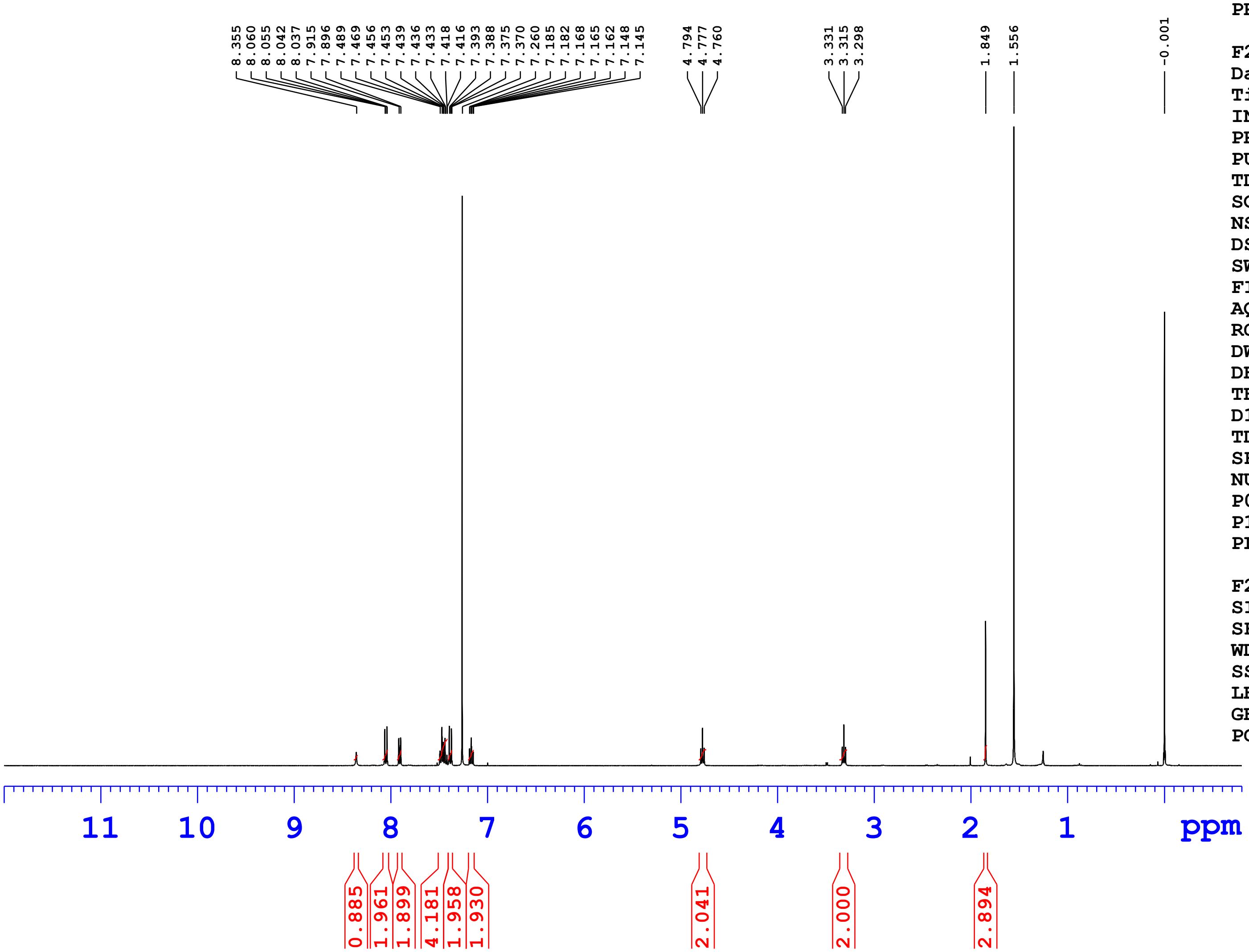


Current Data Parameters

| | |
|---|---|
| NAME | ZINC 13619108 |
| EXPNO | 1 |
| PROCNO | 1 |

F2 - Acquisition Parameters

| | |
|---|---|
| Date_ | 20240916 |
| Time | 11.02 h |
| INSTRUM | Avance Neo Nanobay 400MHz |
| PROBHD | Z163739_0392 ( |
| PULPROG | zg30 |
| TD | 65536 |
| SOLVENT | CDCl3 |
| NS | 16 |
| DS | 0 |
| SWH | 8620.689 Hz |
| FIDRES | 0.263083 Hz |
| AQ | 3.8010881 sec |
| RG | 101 |
| DW | 58.000 usec |
| DE | 13.14 usec |
| TE | 298.1 K |
| D1 | 1.00000000 sec |
| TD0 | 1 |
| SFO1 | 400.3024719 MHz |
| NUC1 | 1H |
| P0 | 2.67 usec |
| P1 | 8.00 usec |
| PLW1 | 23.41300011 W |

F2 - Processing parameters

| | |
|---|---|
| SI | 65536 |
| SF | 400.3000101 MHz |
| WDW | EM |
| SSB | 0 |
| LB | 0.30 Hz |
| GB | 0 |
| PC | 1.00 |

ZINC 13619108
1H CDCl3
16-09-2024

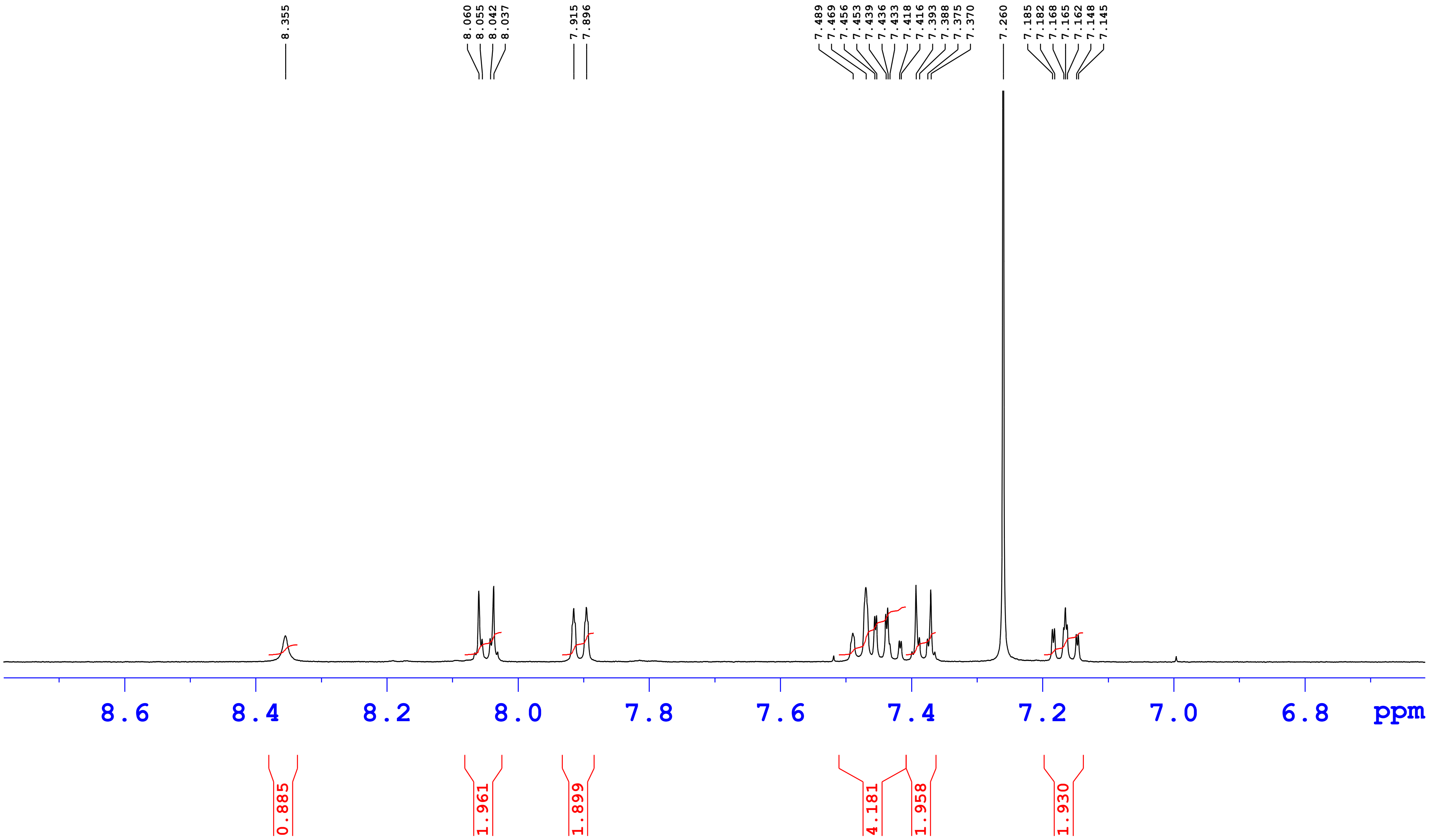

ZINC 13619108
1H CDCl3
16-09-2024

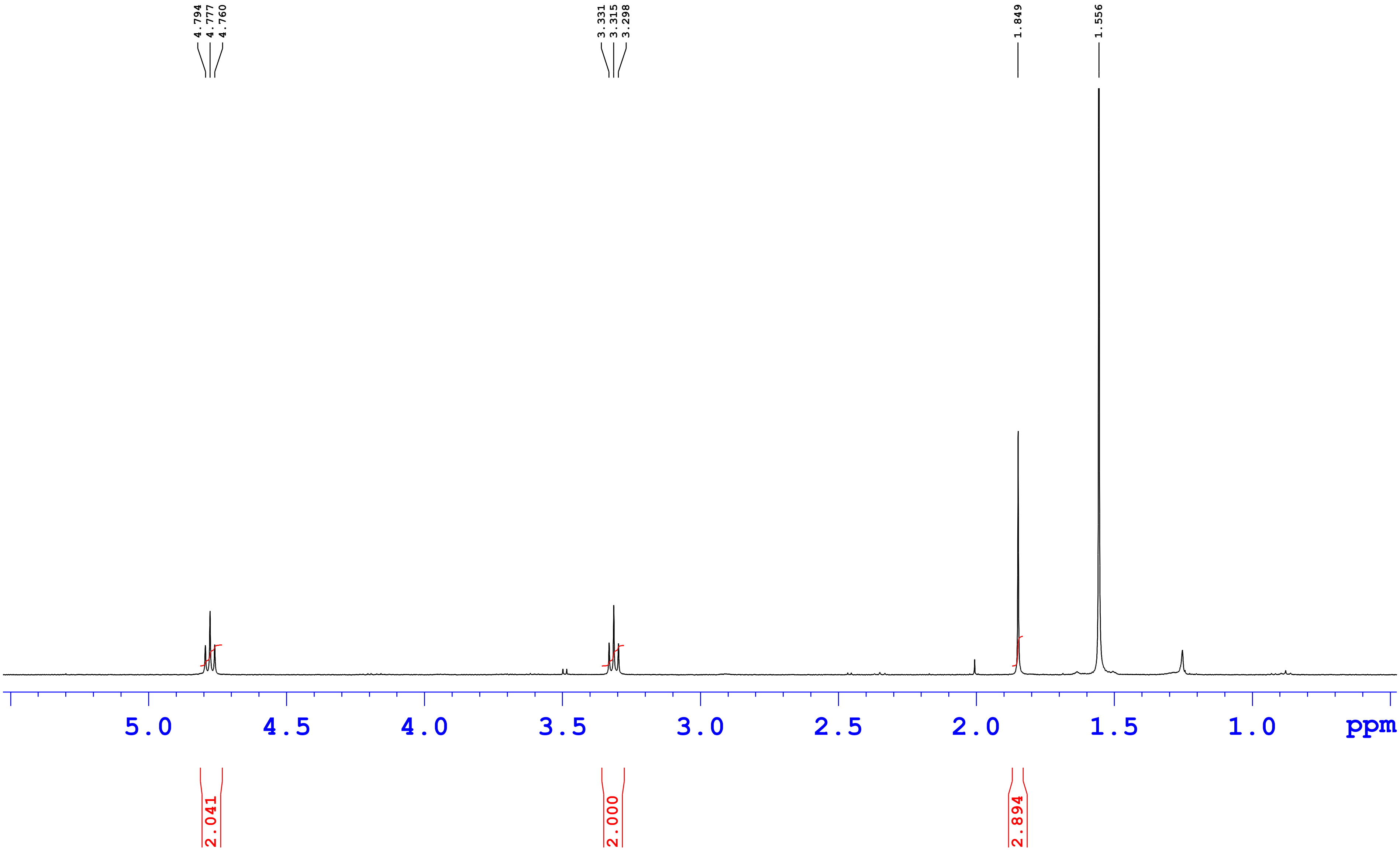

ZINC 13619108

17-Sep-202400:07:11
MPA-0.1%FA In H2O MPB-0.1%FA IN ACN
T/B%-0/5,16/90,20/90,21/5,25/5
Column:Eclipse XDB-C18,4.6*150mm,3.5µm

16092024_054

5: Diode Array
260
Range: 2.084

| Time | Height | Area | Area% |
|---|---|---|---|
| 13.56 | 5850 | 888.90 | 0.31 |
| 14.13 | 2075875 | 281082.06 | 99.46 |
| 14.80 | 5028 | 628.35 | 0.22 |

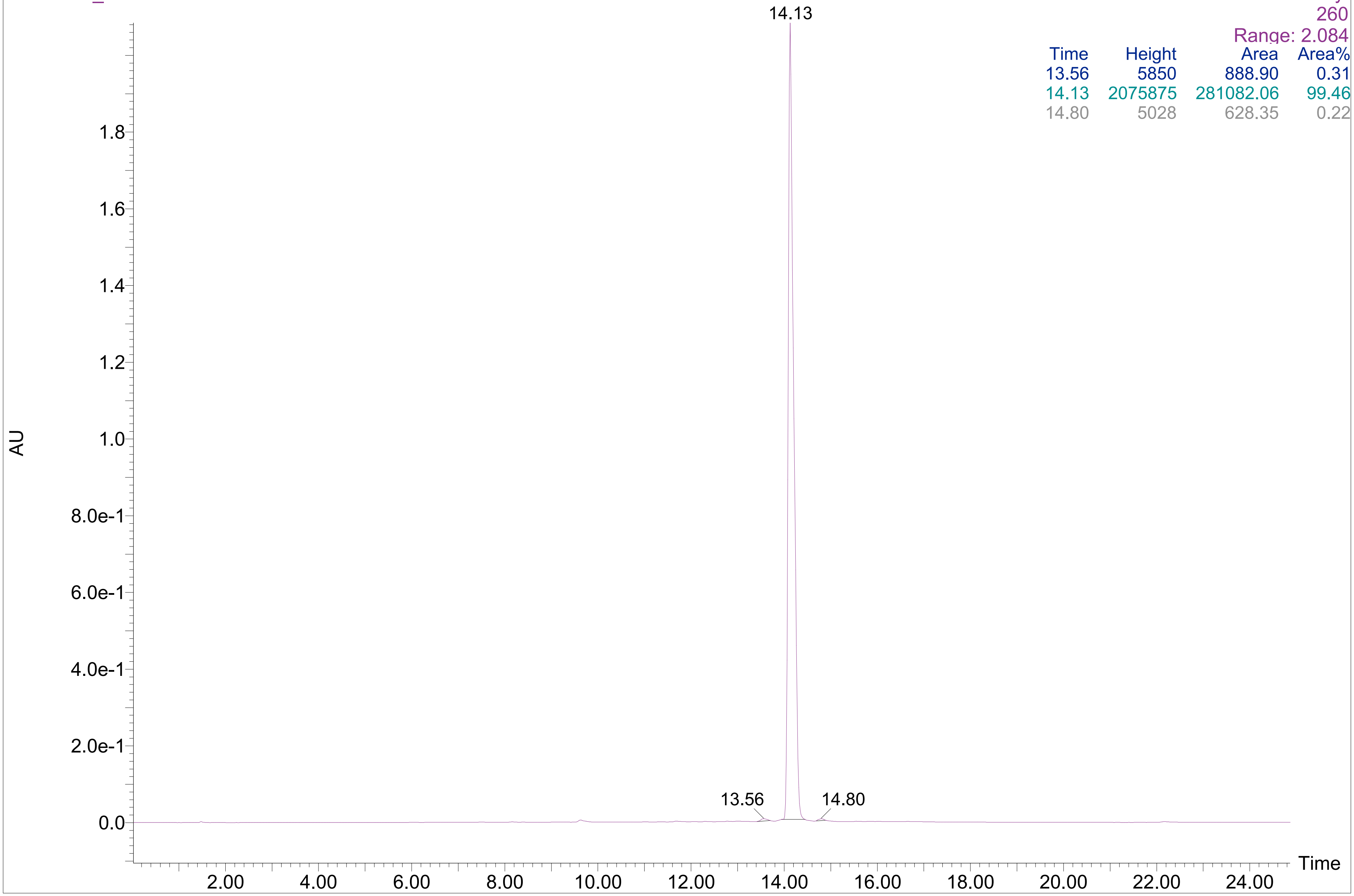

ZINC 13619108

17-Sep-202400:07:11
MPA-0.1%FA In H2O MPB-0.1%FA IN ACN
T/B%-0/5,16/90,20/90,21/5,25/5
Column:Eclipse XDB-C18,4.6*150mm,3.5µm

16092024_054

5: Diode Array
260
Range: 2.084

| Time | Height | Area | Area% |
|---|---|---|---|
| 13.56 | 5850 | 888.90 | 0.31 |
| 14.13 | 2075875 | 281082.06 | 99.46 |
| 14.80 | 5028 | 628.35 | 0.22 |

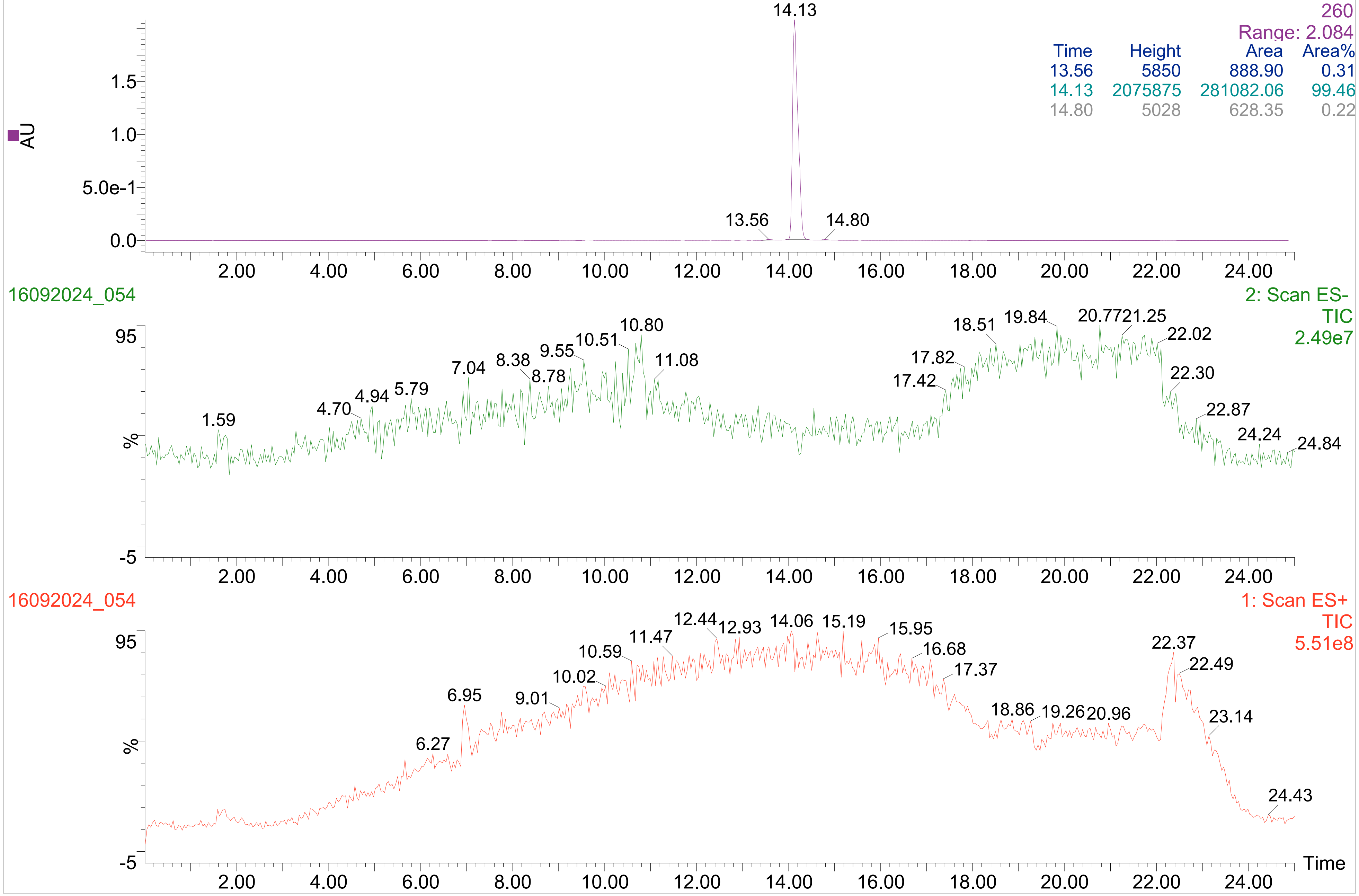

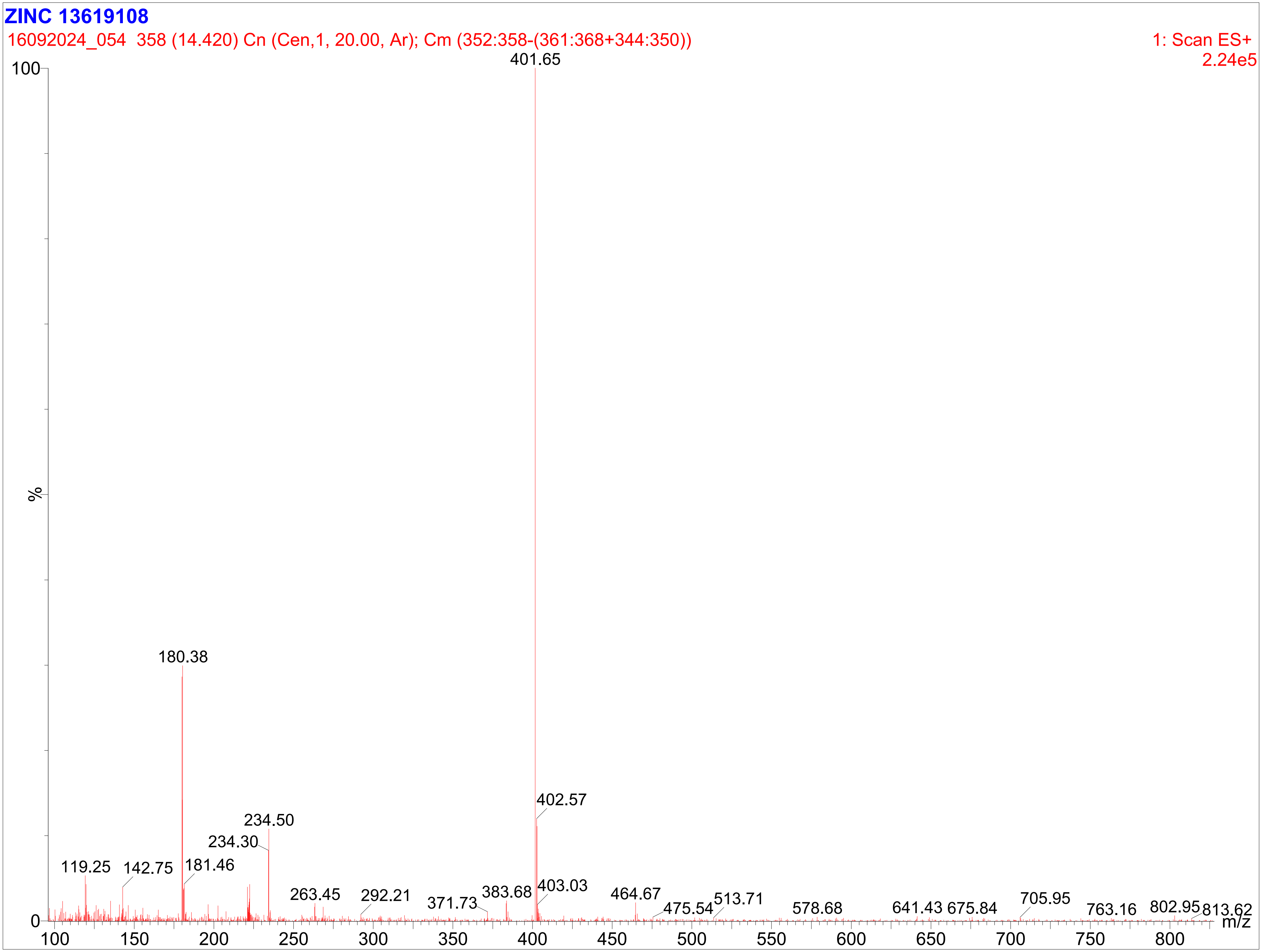

ZINC 13619108
16092024_054 358 (14.420) Cn (Cen,1, 20.00, Ar); Cm (352:358-(361:368+344:350))
1: Scan ES+
2.24e5
401.65
100
%
0
180.38
402.57
234.50
234.30
119.25
142.75
181.46
263.45
292.21
371.73
383.68
403.03
464.67
475.54
513.71
578.68
641.43
675.84
705.95
763.16
802.95
813.62
m/z
100
150
200
250
300
350
400
450
500
550
600
650
700
750
800

ZINC 13619108-A
1H CDCl3
24-09-2024

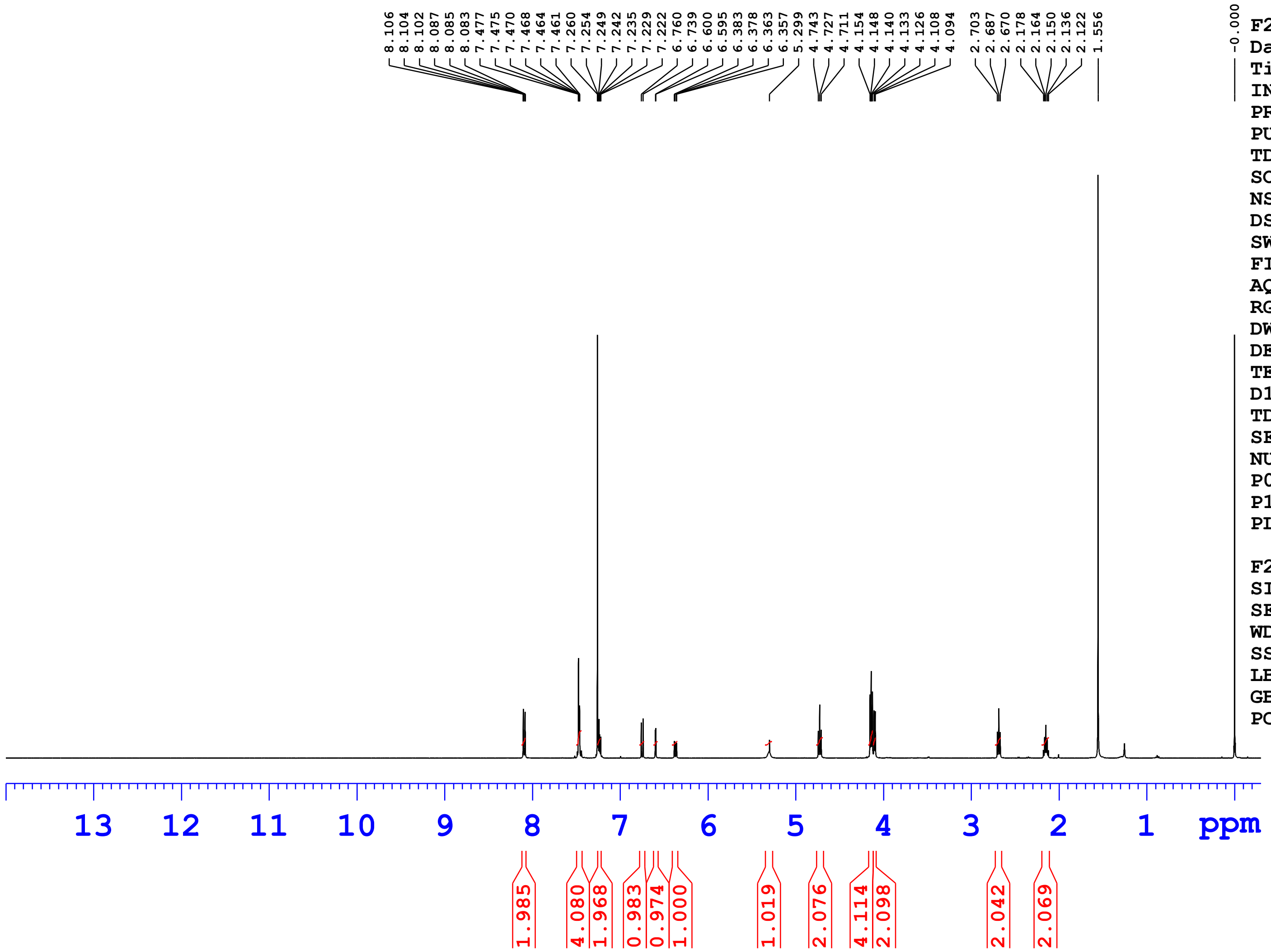


Current Data Parameters
NAME ZINC 13619108-A
EXPNO 1
PROCNO 1

F2 - Acquisition Parameters
Date_ 20240924
Time 18.15 h
INSTRUM Avance Neo Nanobay 400MHz
PROBHD Z163739_0392 (
PULPROG zg30
TD 65536
SOLVENT CDCl3
NS 16
DS 0
SWH 8620.689 Hz
FIDRES 0.263083 Hz
AQ 3.8010881 sec
RG 101
DW 58.000 usec
DE 13.14 usec
TE 298.1 K
D1 1.00000000 sec
TD0 1
SFO1 400.3024719 MHz
NUC1 1H
P0 2.67 usec
P1 8.00 usec
PLW1 23.41300011 W

F2 - Processing parameters
SI 65536
SF 400.3000102 MHz
WDW EM
SSB 0
LB 0.30 Hz
GB 0
PC 1.00

ZINC 13619108-A
1H CDCl3
24-09-2024

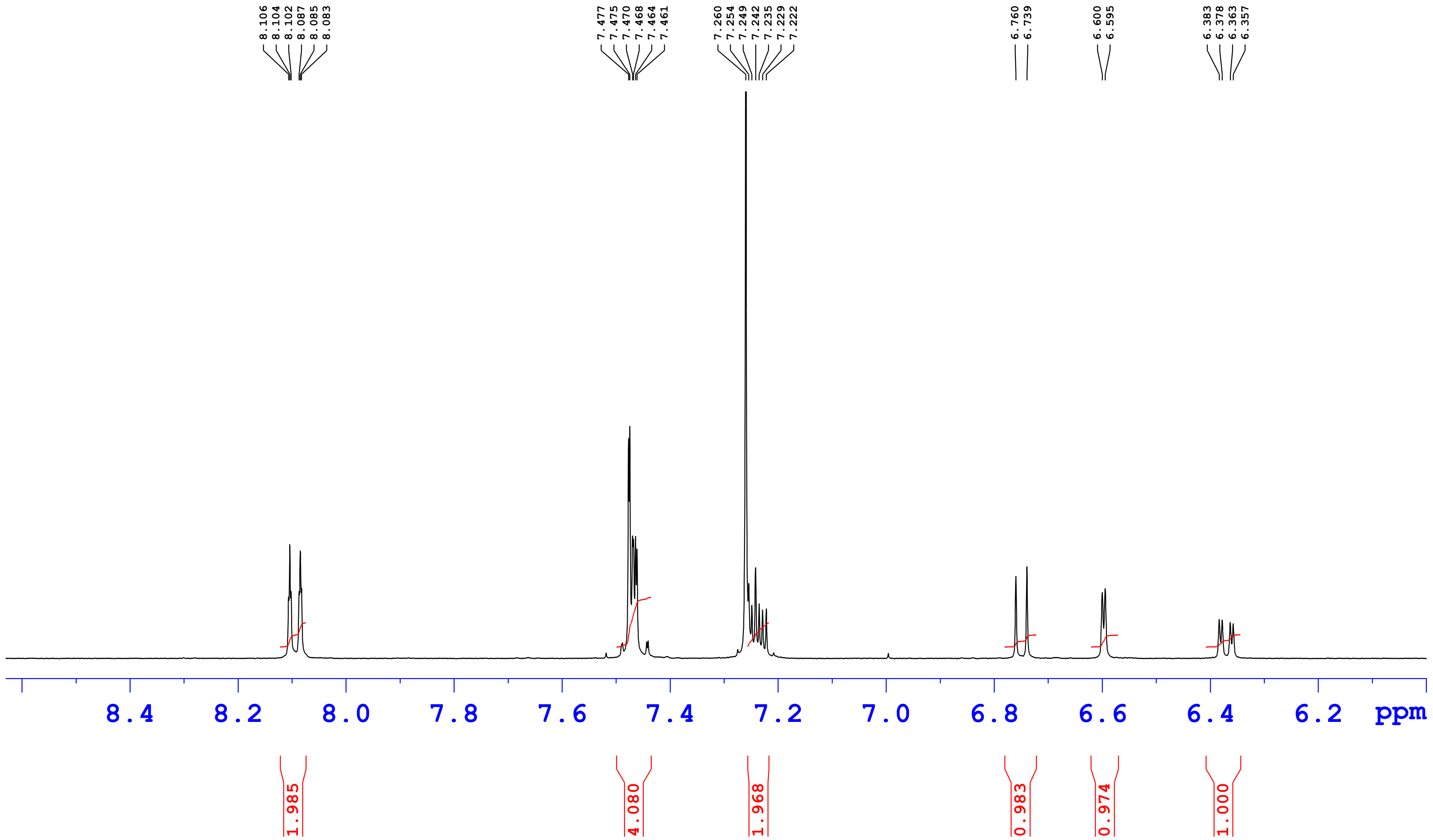

ZINC 13619108-A
1H CDCl3
24-09-2024

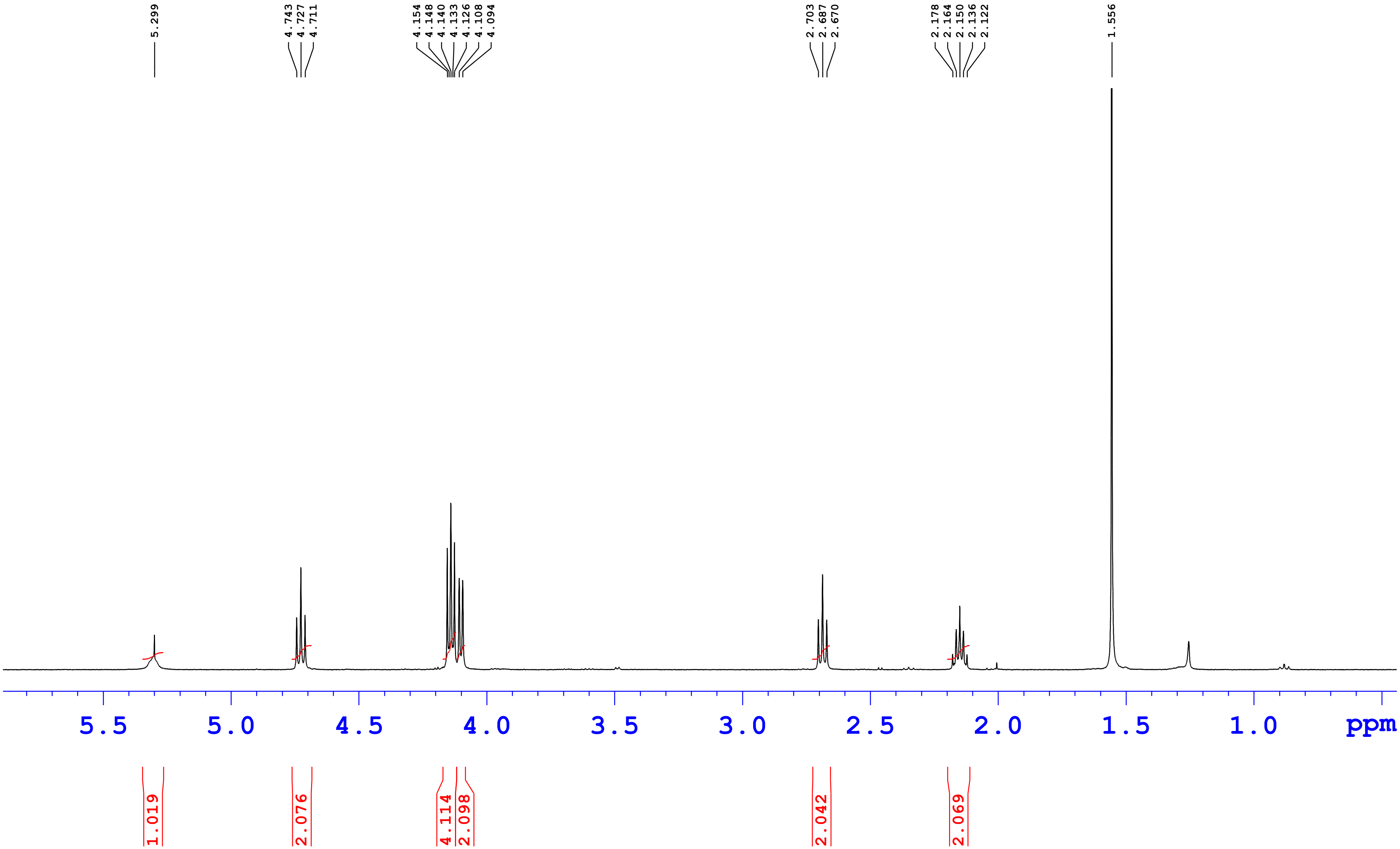

**ZINC13619108-A**

**25-Sep-202421:32:48**
**MPA-0.1%FA in H2O MPB-ACN**
**T/B%-0/5,16/90,20/90,21/5,25/5**
**InertClone ODS(2),4.6*150mm,5µm**

25092024_056

5: Diode Array
235.869
Range: 2.615

| Time | Height | Area | Area% |
|---|---|---|---|
| 13.34 | 2548041 | 393321.84 | 98.81 |
| 14.08 | 12942 | 1481.67 | 0.37 |
| 14.63 | 21370 | 2225.74 | 0.56 |
| 15.36 | 9718 | 1043.27 | 0.26 |

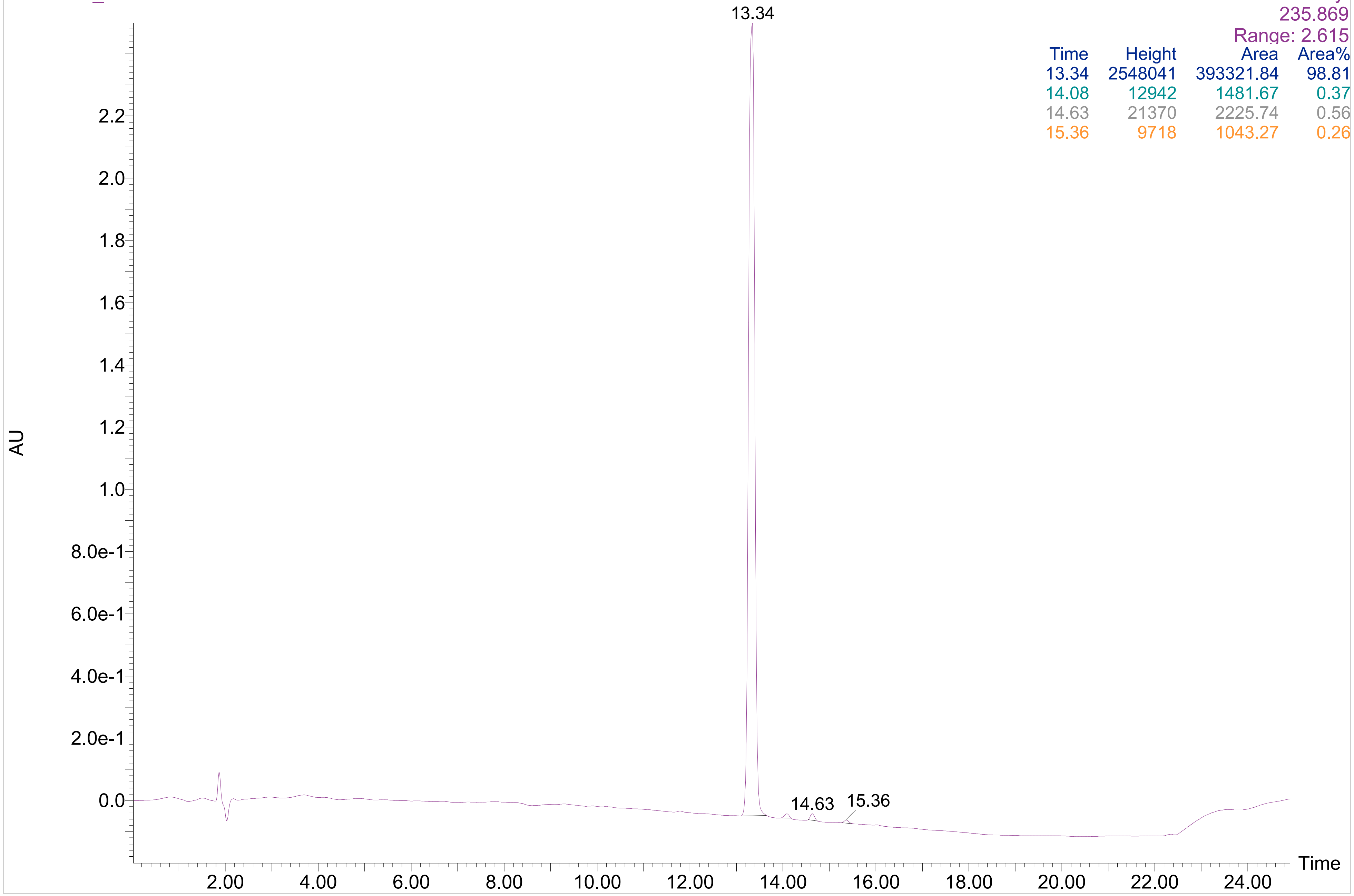

**ZINC13619108-A**

**25-Sep-202421:32:48**
**MPA-0.1%FA in H2O MPB-ACN**
**T/B%-0/5,16/90,20/90,21/5,25/5**
**InertClone ODS(2),4.6*150mm,5μm**

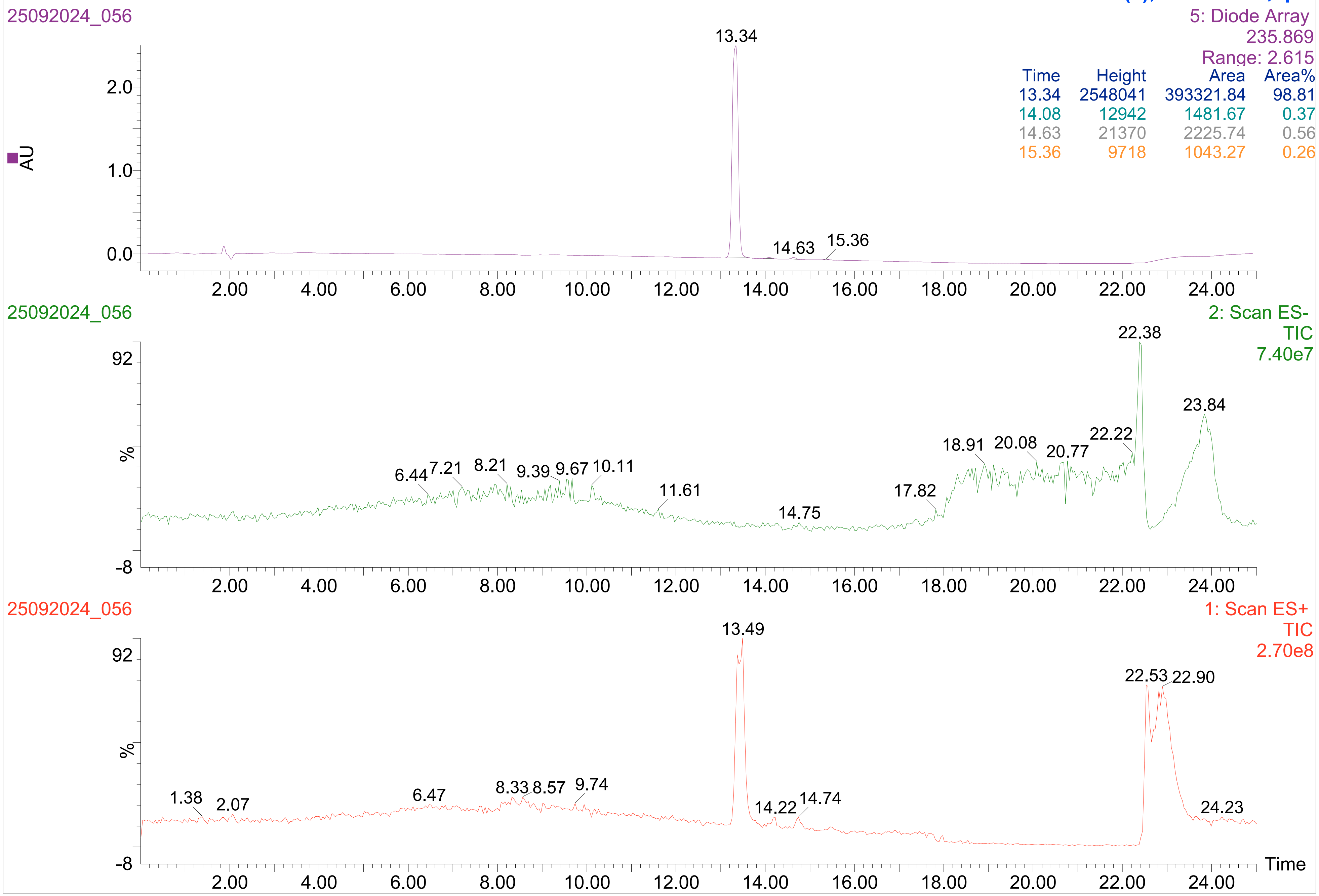


| Time | Height | Area | Area% |
|---|---|---|---|
| 13.34 | 2548041 | 393321.84 | 98.81 |
| 14.08 | 12942 | 1481.67 | 0.37 |
| 14.63 | 21370 | 2225.74 | 0.56 |
| 15.36 | 9718 | 1043.27 | 0.26 |

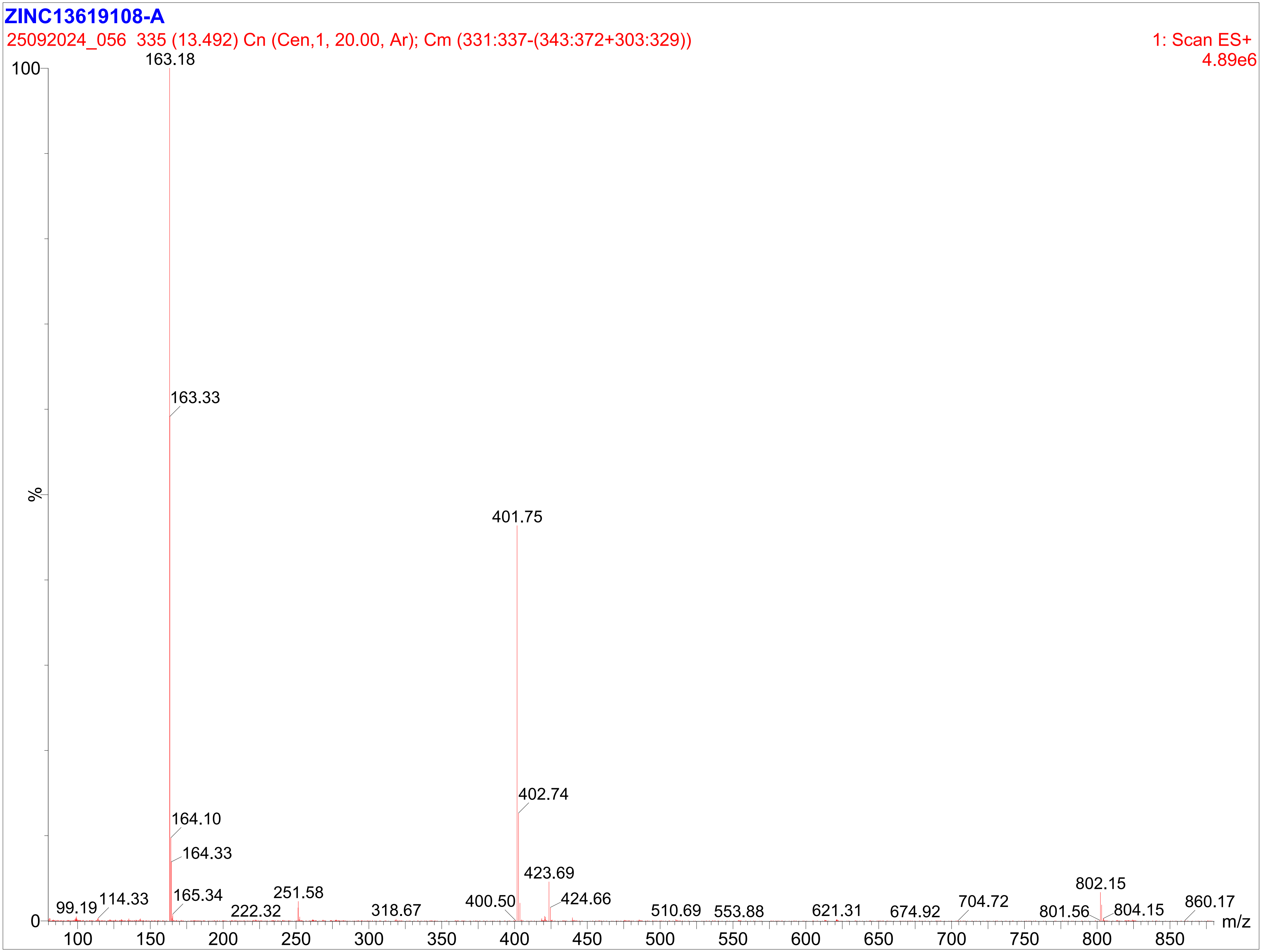
ZINC13619108-A
25092024_056 335 (13.492) Cn (Cen,1, 20.00, Ar); Cm (331:337-(343:372+303:329))
1: Scan ES+
4.89e6
%
m/z
163.18
163.33
164.10
164.33
165.34
401.75
402.74
423.69
424.66
99.19
114.33
222.32
251.58
318.67
400.50
510.69
553.88
621.31
674.92
704.72
801.56
802.15
804.15
860.17

ZINC 15938334-A
1H DMSO-D6
24-09-2024

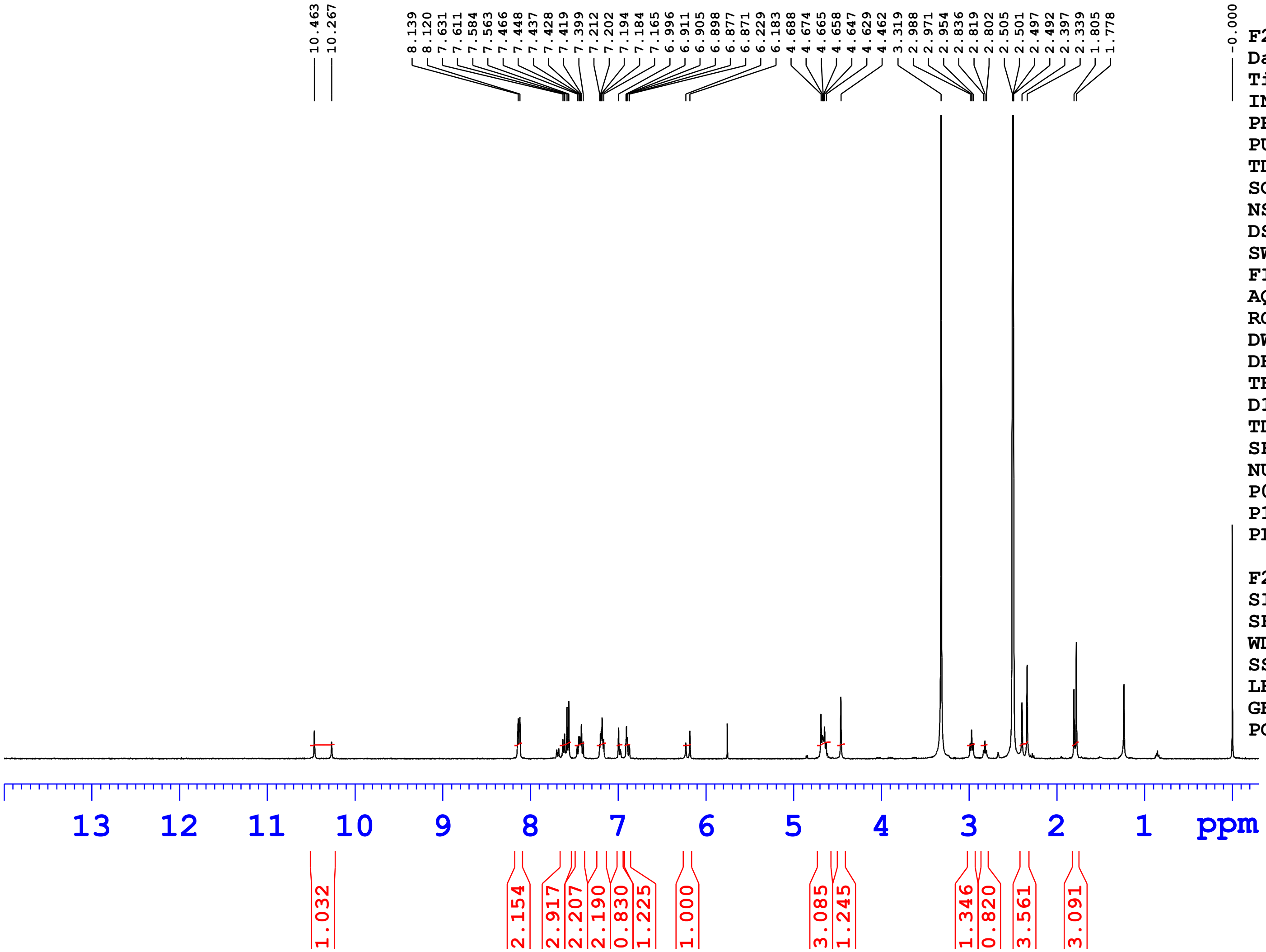


Current Data Parameters

| | |
|---|---|
| NAME | ZINC 15938334-A |
| EXPNO | 1 |
| PROCNO | 1 |

F2 - Acquisition Parameters

| | |
|---|---|
| Date_ | 20240924 |
| Time | 10.57 h |
| INSTRUM | Avance Neo Nanobay 400MHz |
| PROBHD | Z163739_0392 ( |
| PULPROG | zg30 |
| TD | 65536 |
| SOLVENT | DMSO |
| NS | 16 |
| DS | 0 |
| SWH | 8620.689 Hz |
| FIDRES | 0.263083 Hz |
| AQ | 3.8010881 sec |
| RG | 101 |
| DW | 58.000 usec |
| DE | 13.14 usec |
| TE | 298.1 K |
| D1 | 1.00000000 sec |
| TD0 | 1 |
| SFO1 | 400.3024719 MHz |
| NUC1 | 1H |
| P0 | 2.67 usec |
| P1 | 8.00 usec |
| PLW1 | 23.41300011 W |

F2 - Processing parameters

| | |
|---|---|
| SI | 65536 |
| SF | 400.3000021 MHz |
| WDW | EM |
| SSB | 0 |
| LB | 0.30 Hz |
| GB | 0 |
| PC | 1.00 |

ZINC 15938334-A
1H DMSO-D6
24-09-2024

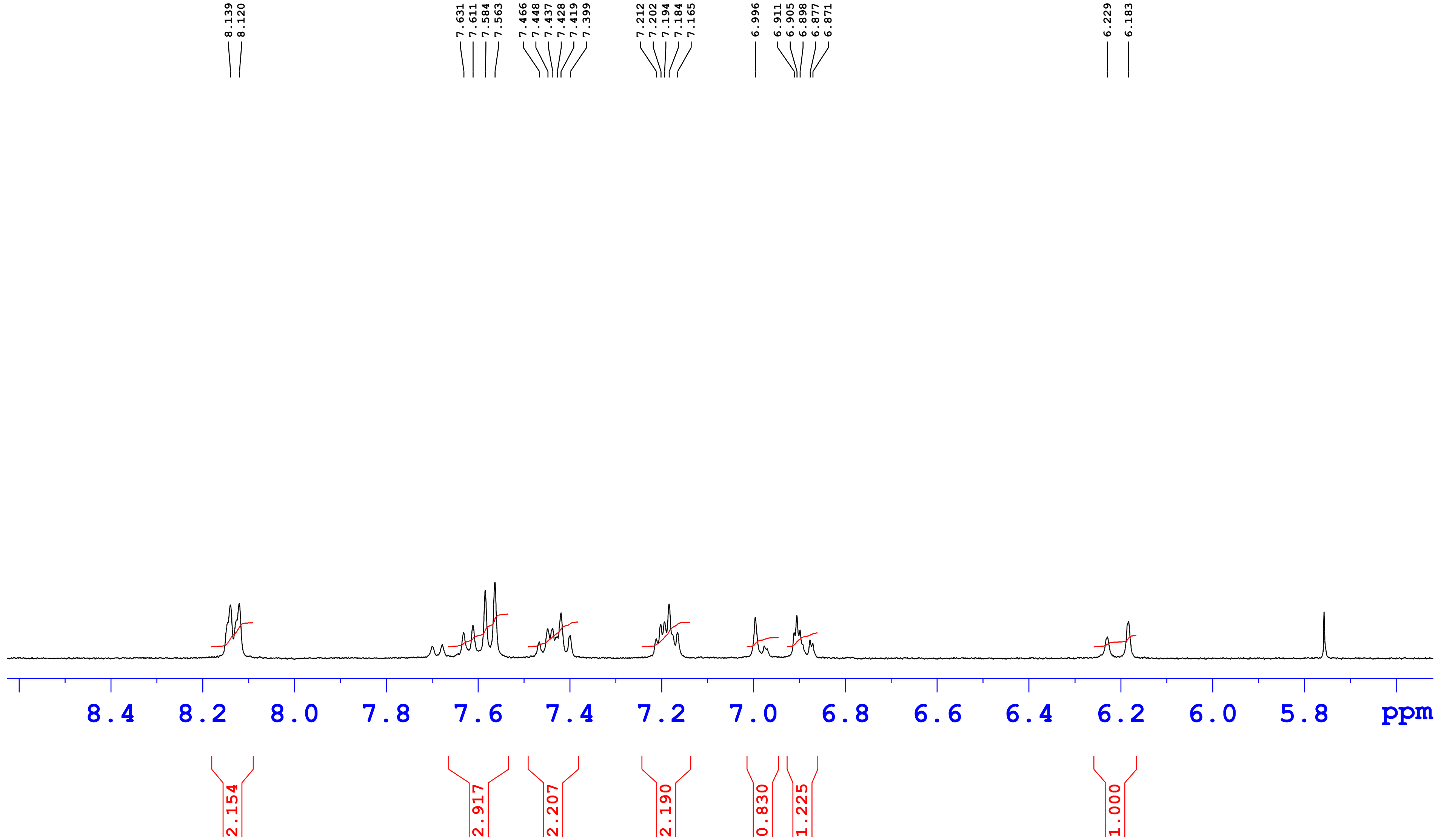

ZINC 15938334-A
1H DMSO-D6
24-09-2024

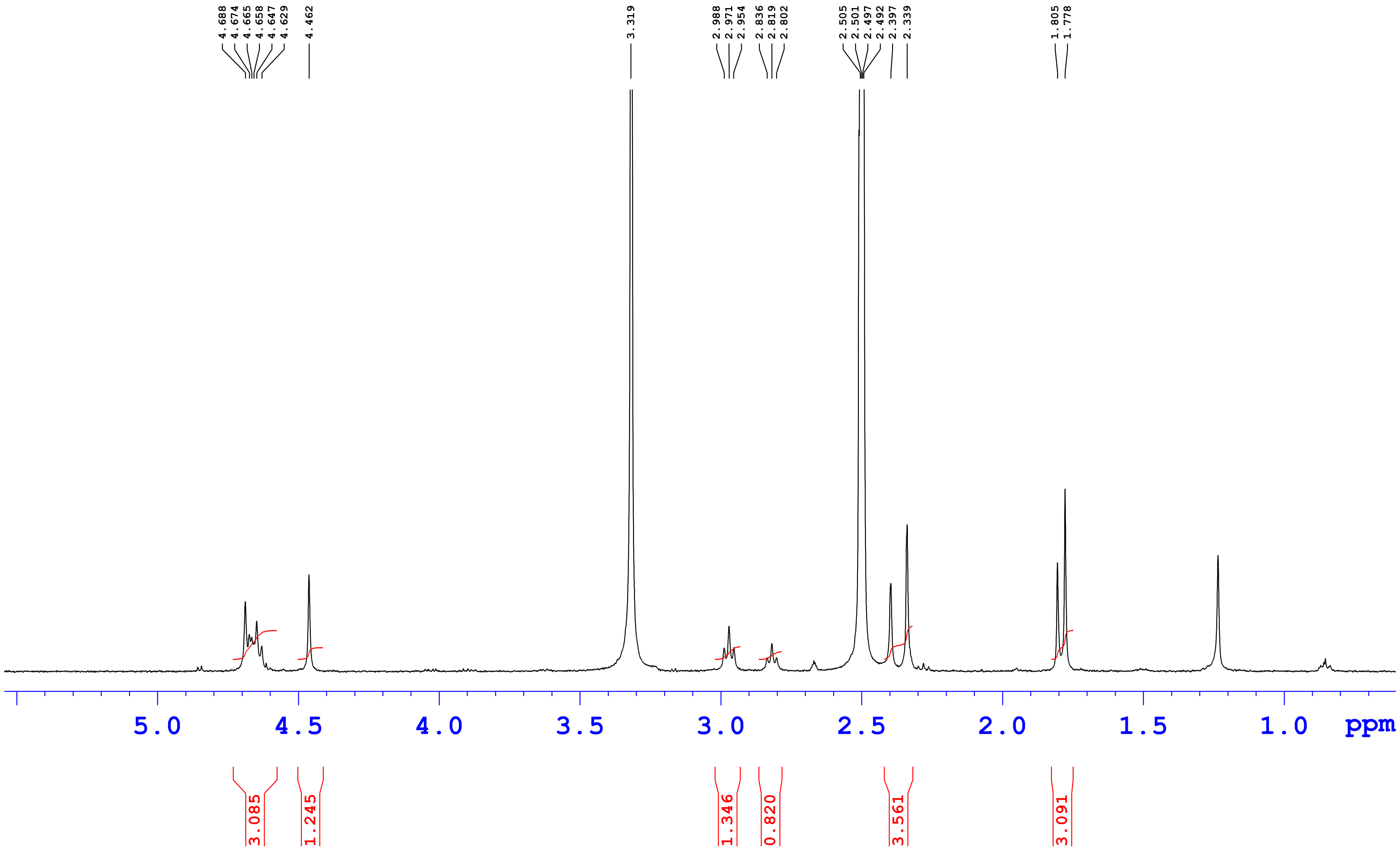

ZINC15938334
1H DMSO-D6
12-09-2024

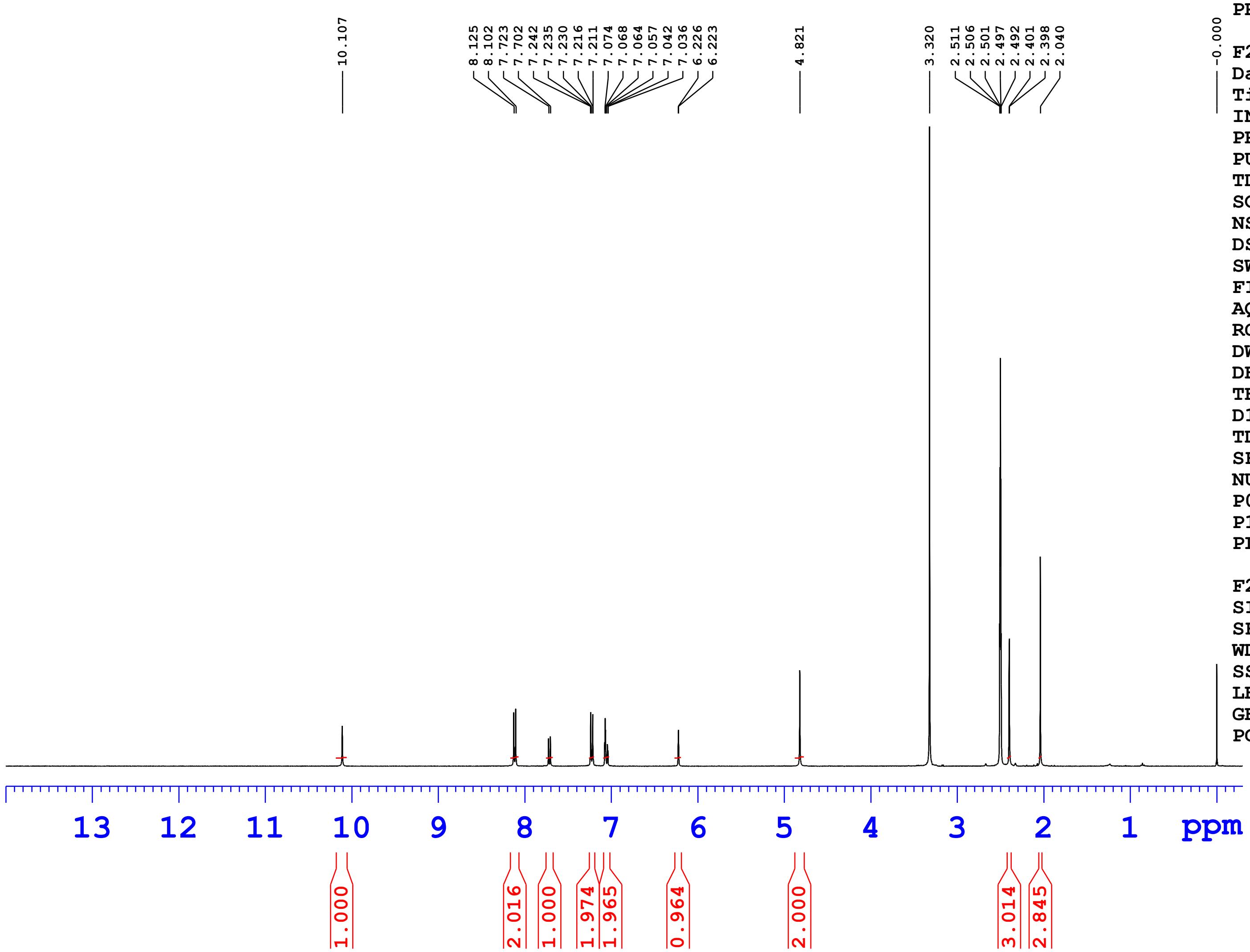


Current Data Parameters

| | |
|---|---|
| NAME | ZINC15938334 |
| EXPNO | 1 |
| PROCNO | 1 |

F2 - Acquisition Parameters

| | |
|---|---|
| Date_ | 20240912 |
| Time | 10.05 h |
| INSTRUM | Avance Neo Nanobay 400MHz |
| PROBHD | Z163739_0392 ( |
| PULPROG | zg30 |
| TD | 65536 |
| SOLVENT | DMSO |
| NS | 16 |
| DS | 0 |
| SWH | 8620.689 Hz |
| FIDRES | 0.263083 Hz |
| AQ | 3.8010881 sec |
| RG | 101 |
| DW | 58.000 usec |
| DE | 13.14 usec |
| TE | 298.2 K |
| D1 | 1.00000000 sec |
| TD0 | 1 |
| SFO1 | 400.3024719 MHz |
| NUC1 | 1H |
| P0 | 2.67 usec |
| P1 | 8.00 usec |
| PLW1 | 23.41300011 W |

F2 - Processing parameters

| | |
|---|---|
| SI | 65536 |
| SF | 400.3000018 MHz |
| WDW | EM |
| SSB | 0 |
| LB | 0.30 Hz |
| GB | 0 |
| PC | 1.00 |

ZINC15938334
1H DMSO-D6
12-09-2024

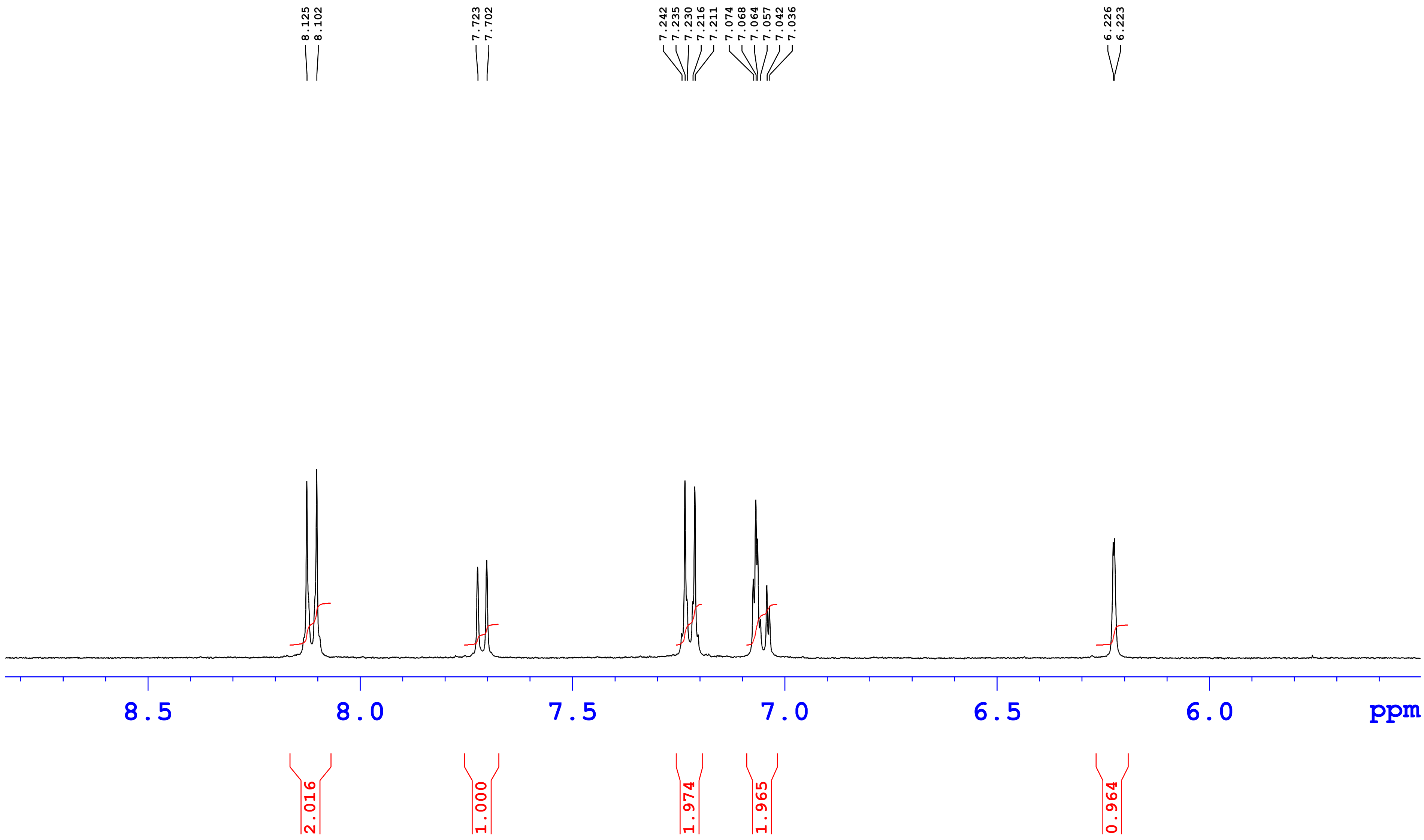

ZINC15938334
1H DMSO-D6
12-09-2024

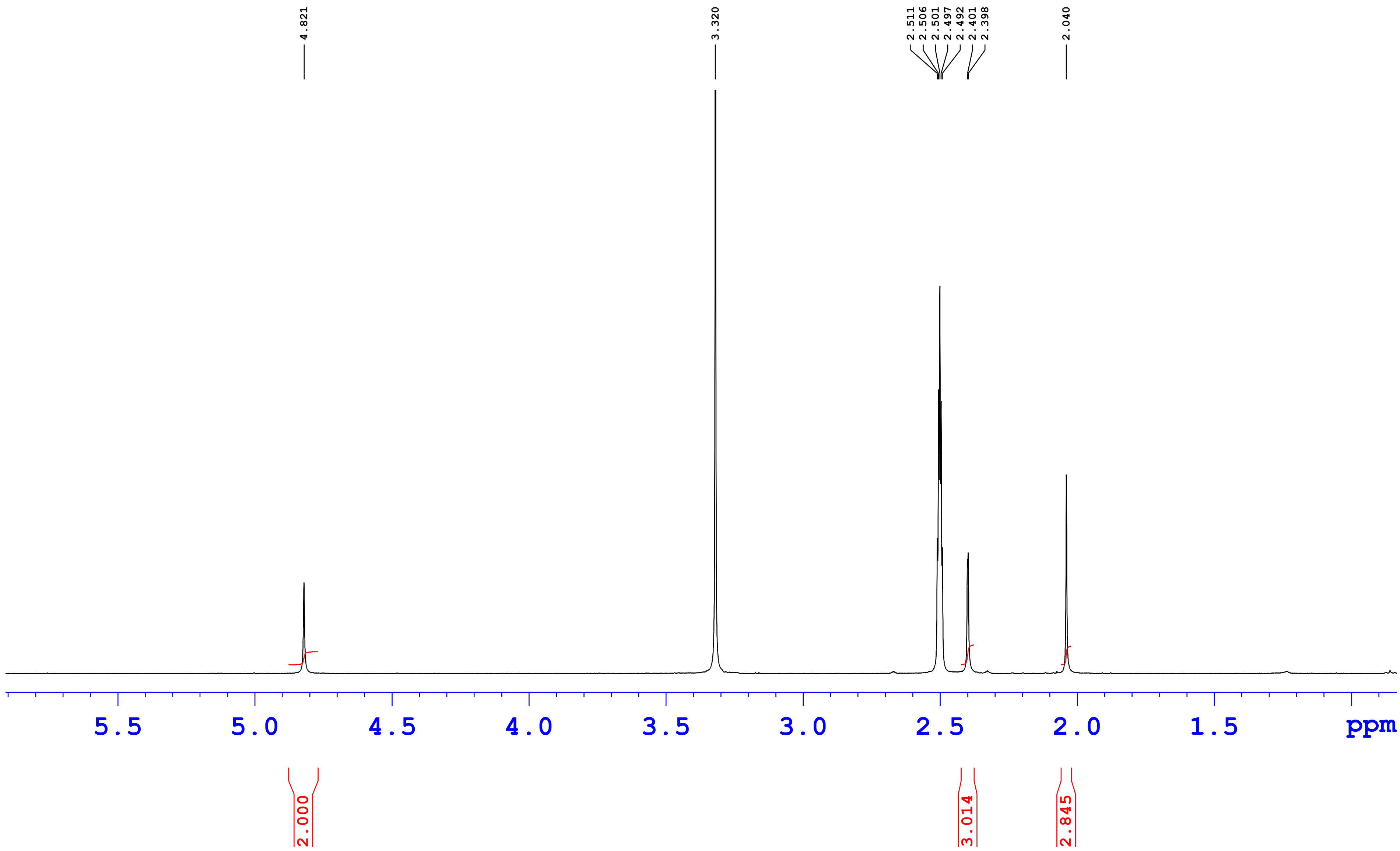

**ZINC15938334**

**12-Sep-202422:10:10**
**Mobile Phase (A)0.1% FA In H2O(B)0.1% FA In ACN**
**Column Eclipse XDB C18 4.6*150mm 3.5µm**
**Gradient T/% B 0/5,16/90,20/90,21/5,25/5**

12092024_136

5: Diode Array
310
Range: 1.069

| Time | Height | Area | Area% |
|---|---|---|---|
| 4.20 | 1851 | 273.00 | 0.19 |
| 7.54 | 7722 | 882.84 | 0.63 |
| 8.67 | 2599 | 325.22 | 0.23 |
| 11.00 | 739 | 107.14 | 0.08 |
| 13.20 | 1067190 | 133645.13 | 95.39 |
| 14.12 | 31032 | 4867.15 | 3.47 |

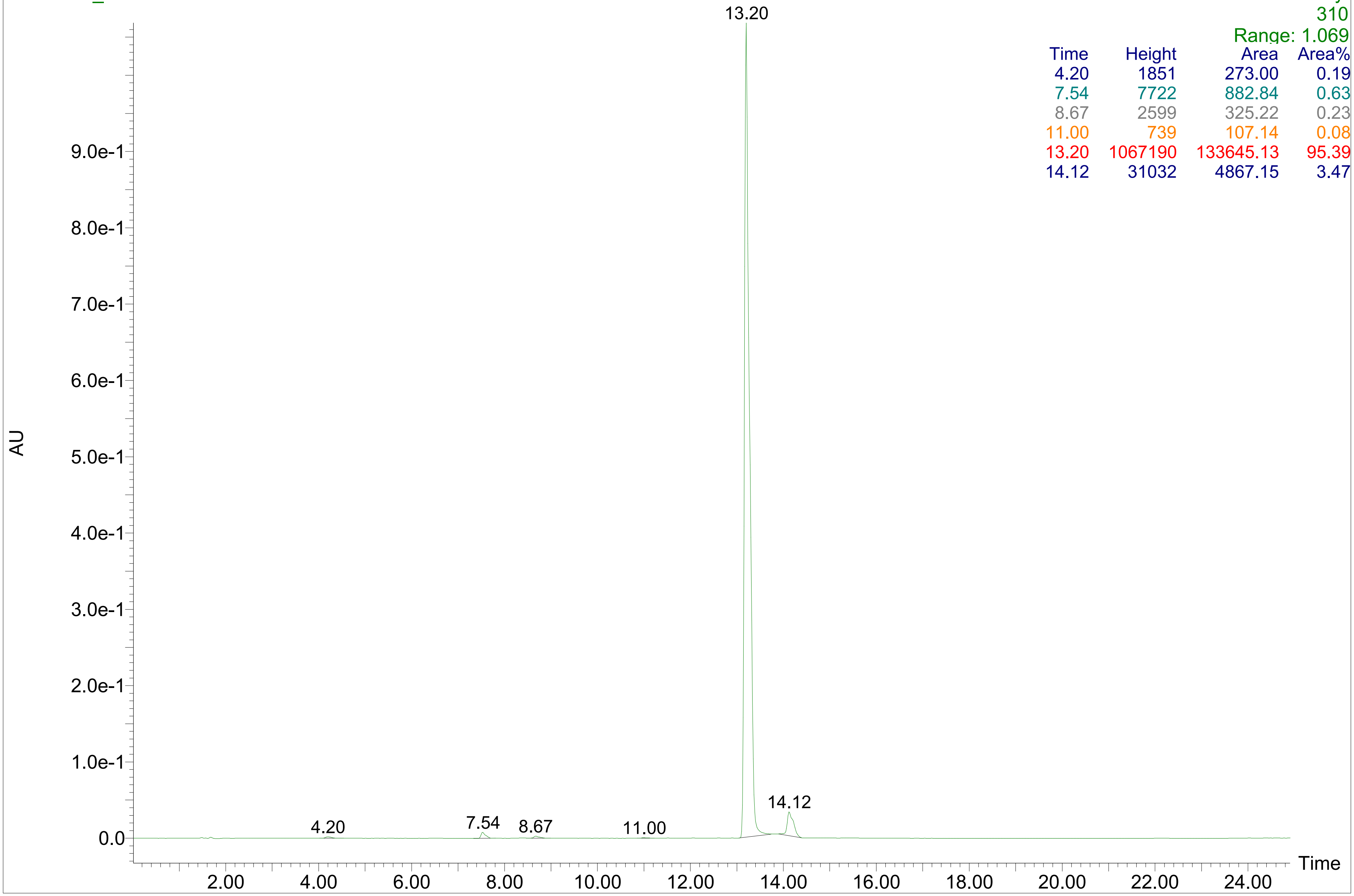

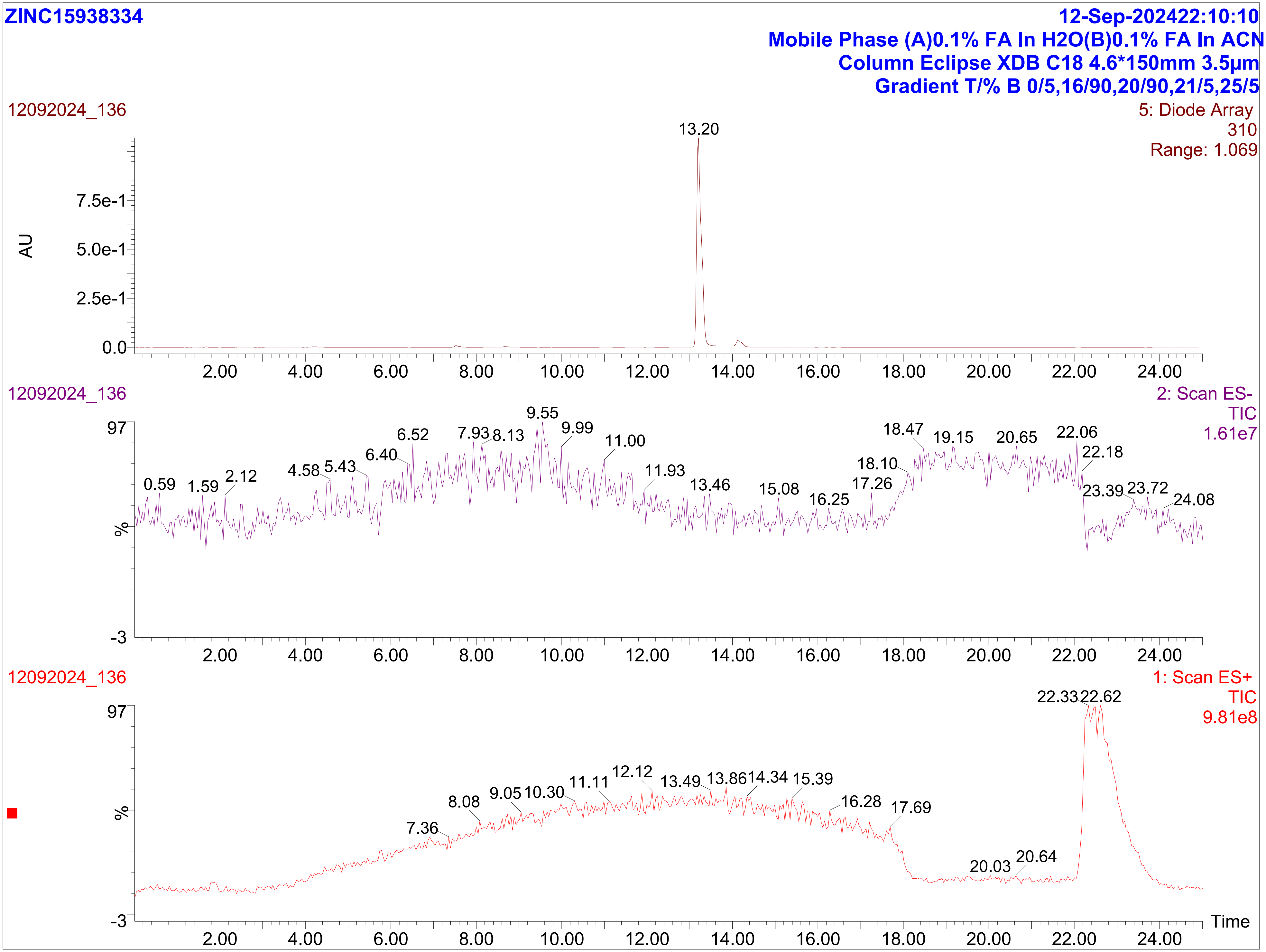

ZINC15938334
12-Sep-202422:10:10
Mobile Phase (A)0.1% FA In H2O(B)0.1% FA In ACN
Column Eclipse XDB C18 4.6*150mm 3.5µm
Gradient T/% B 0/5,16/90,20/90,21/5,25/5
12092024_136
5: Diode Array
310
Range: 1.069
13.20
AU
12092024_136
2: Scan ES-
TIC
1.61e7
12092024_136
1: Scan ES+
TIC
9.81e8
22.33
22.62
Time

ZINC15938334

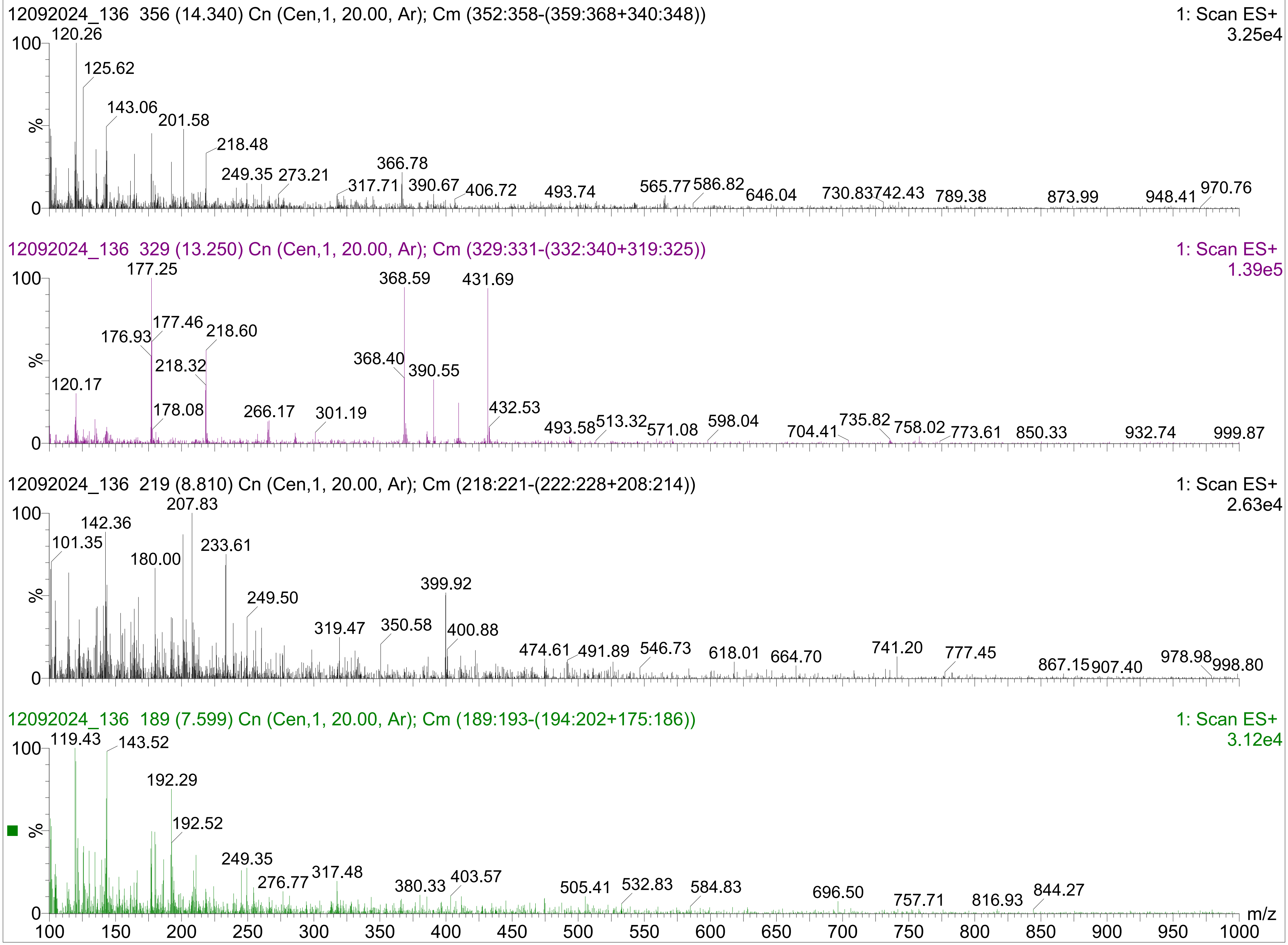

ZINC24080467
1H CDCl3
23-09-2024

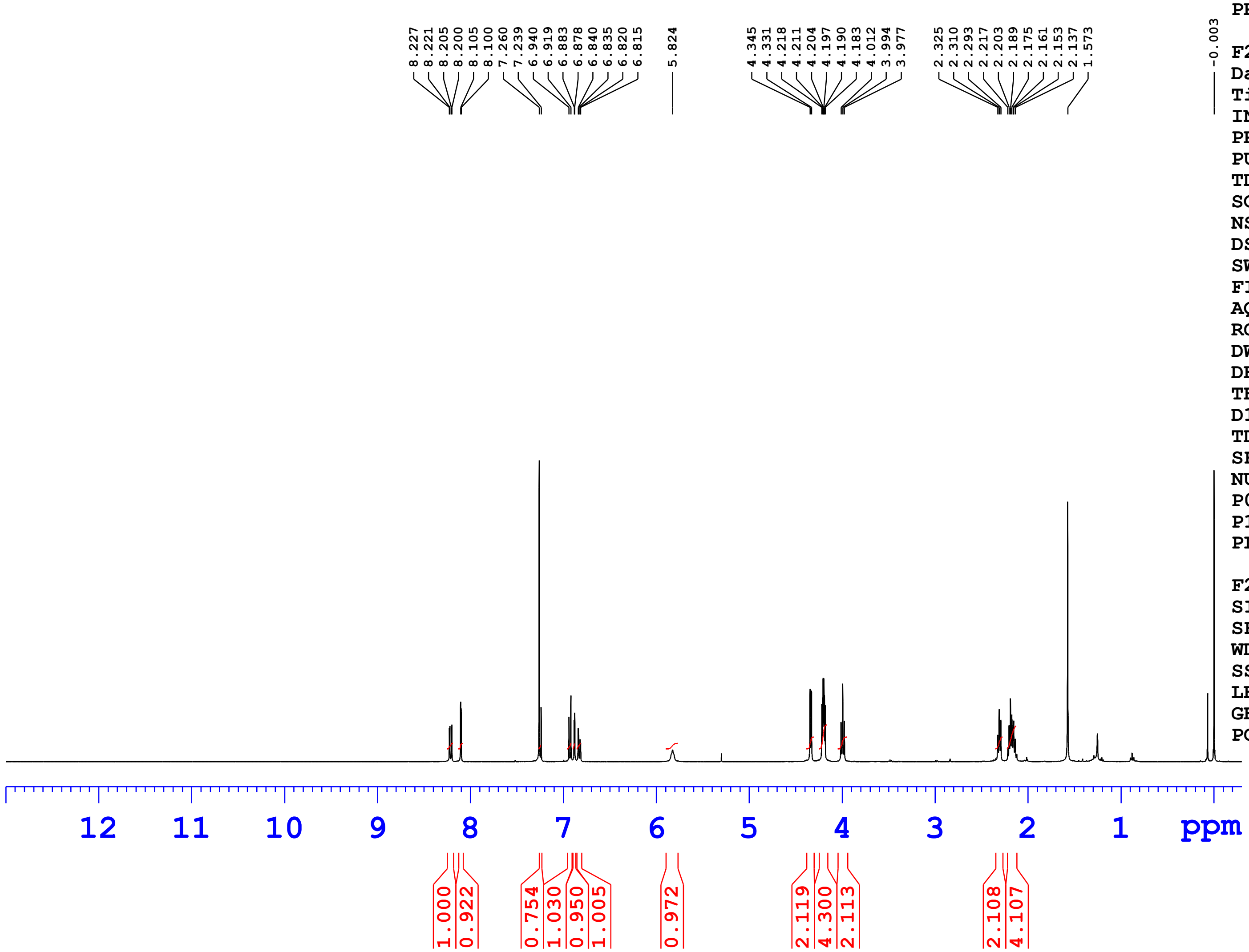


Current Data Parameters
NAME ZINC24080467
EXPNO 1
PROCNO 1

F2 - Acquisition Parameters
Date_ 20240923
Time 17.40 h
INSTRUM Avance Neo Nanobay 400MHz
PROBHD Z163739_0392 (
PULPROG zg30
TD 65536
SOLVENT CDCl3
NS 16
DS 0
SWH 8620.689 Hz
FIDRES 0.263083 Hz
AQ 3.8010881 sec
RG 101
DW 58.000 usec
DE 13.14 usec
TE 298.1 K
D1 1.00000000 sec
TD0 1
SFO1 400.3024719 MHz
NUC1 1H
P0 2.67 usec
P1 8.00 usec
PLW1 23.41300011 W

F2 - Processing parameters
SI 65536
SF 400.3000102 MHz
WDW EM
SSB 0
LB 0.30 Hz
GB 0
PC 1.00

ZINC24080467
1H CDCl3
23-09-2024

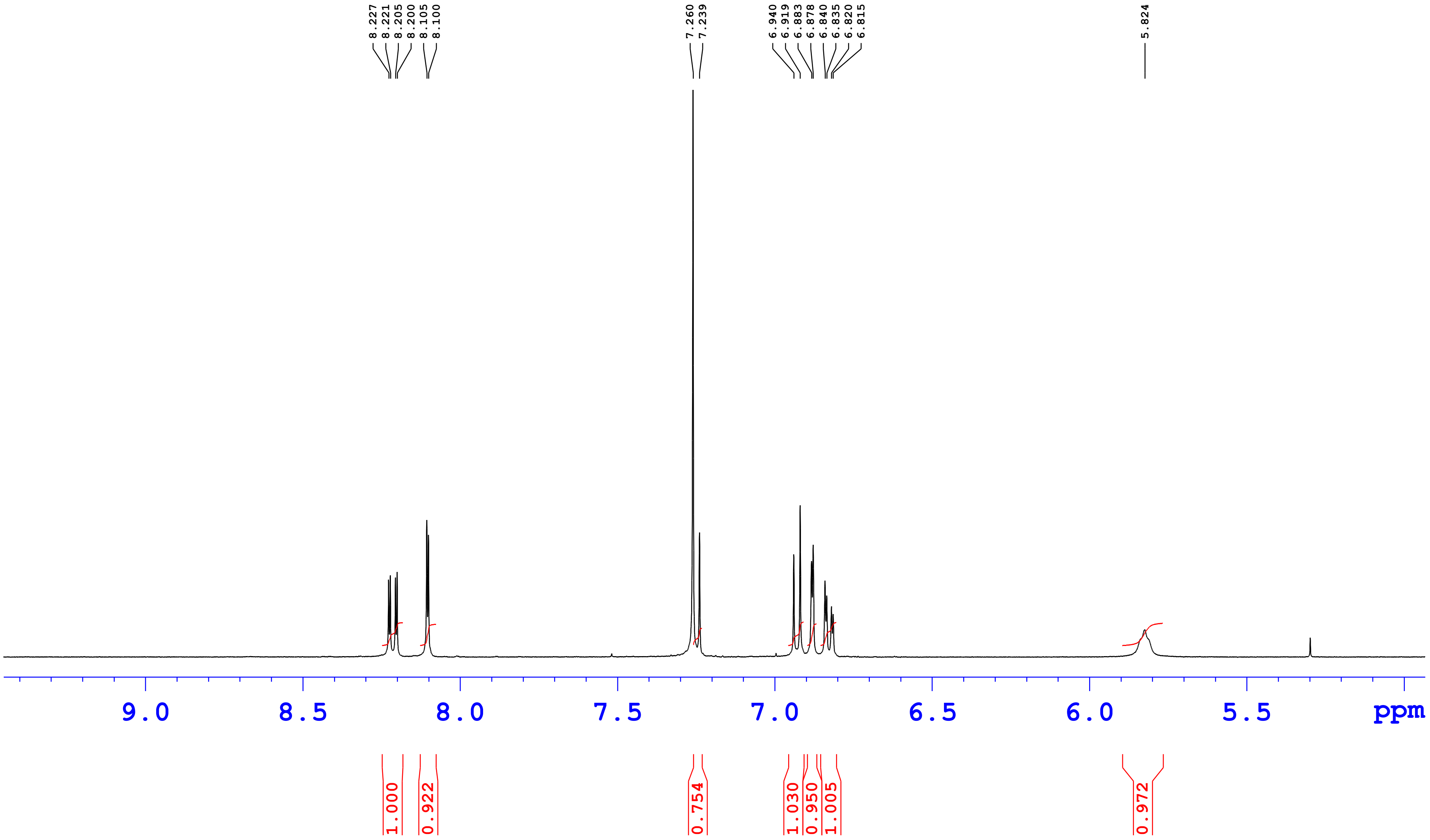

ZINC24080467
1H CDCl3
23-09-2024

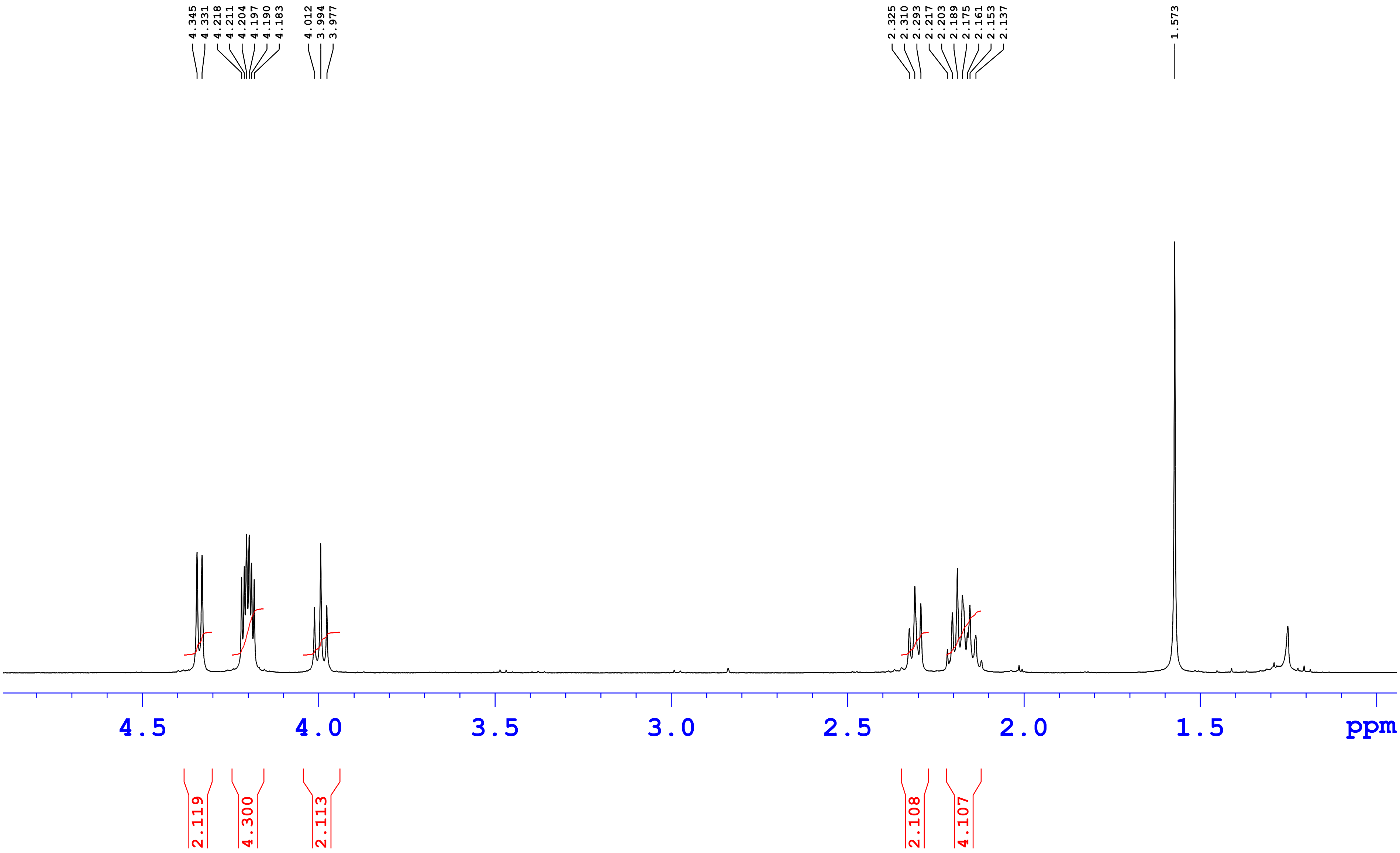

ZINC24080467

24-Sep-202401:03:08
MPA(A) 0.1% FA In H2O(B) 0.1% FA In ACN
Gradient T/%B 0/5,16/90,20/90,21/5,25/5
Column:Eclipse XDB-C18,4.6*150mm,3.5µm

23092024_062

5: Diode Array
320
Range: 2.376

| Time | Height | Area | Area% |
|---|---|---|---|
| 8.24 | 3624 | 271.93 | 0.07 |
| 8.42 | 7919 | 651.81 | 0.17 |
| 8.56 | 14198 | 1529.75 | 0.40 |
| 10.62 | 2374322 | 372407.25 | 98.16 |
| 11.76 | 36633 | 4513.25 | 1.19 |

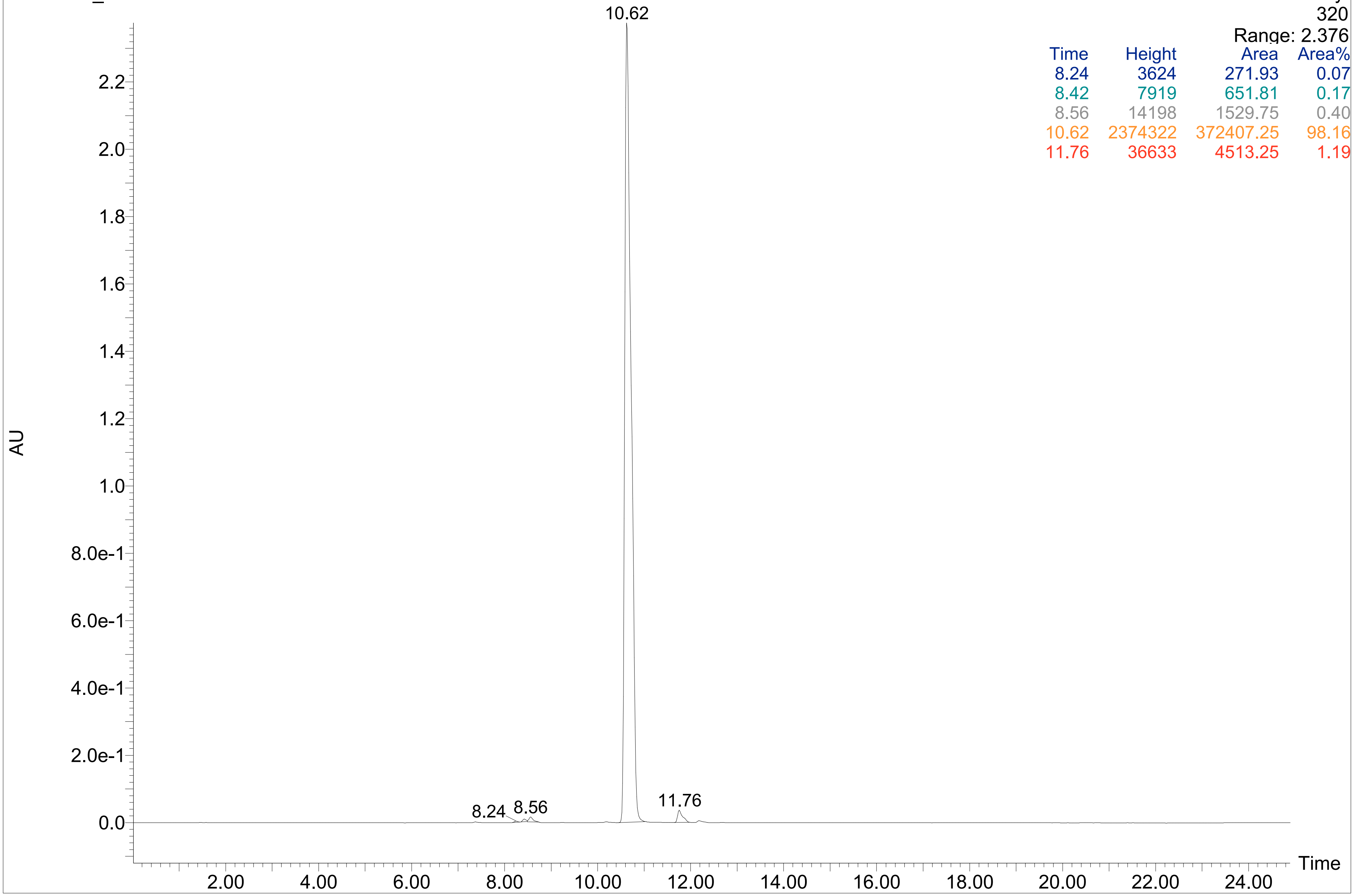

**ZINC24080467**

**24-Sep-202401:03:08**
**MPA(A) 0.1% FA In H2O(B) 0.1% FA In ACN**
**Gradient T/%B 0/5,16/90,20/90,21/5,25/5**
**Column:Eclipse XDB-C18,4.6*150mm,3.5µm**

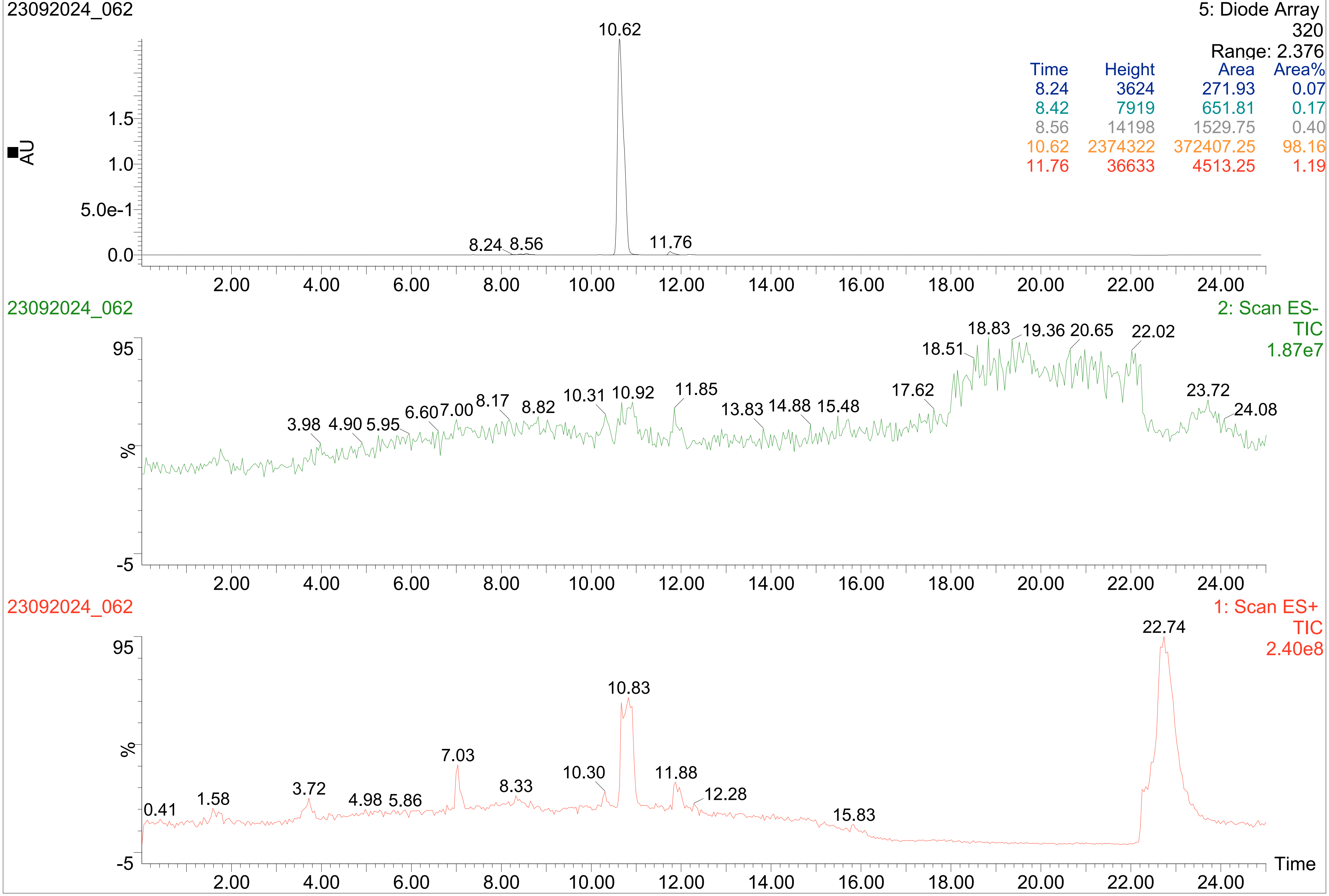


| Time | Height | Area | Area% |
|---|---|---|---|
| 8.24 | 3624 | 271.93 | 0.07 |
| 8.42 | 7919 | 651.81 | 0.17 |
| 8.56 | 14198 | 1529.75 | 0.40 |
| 10.62 | 2374322 | 372407.25 | 98.16 |
| 11.76 | 36633 | 4513.25 | 1.19 |

ZINC24080467

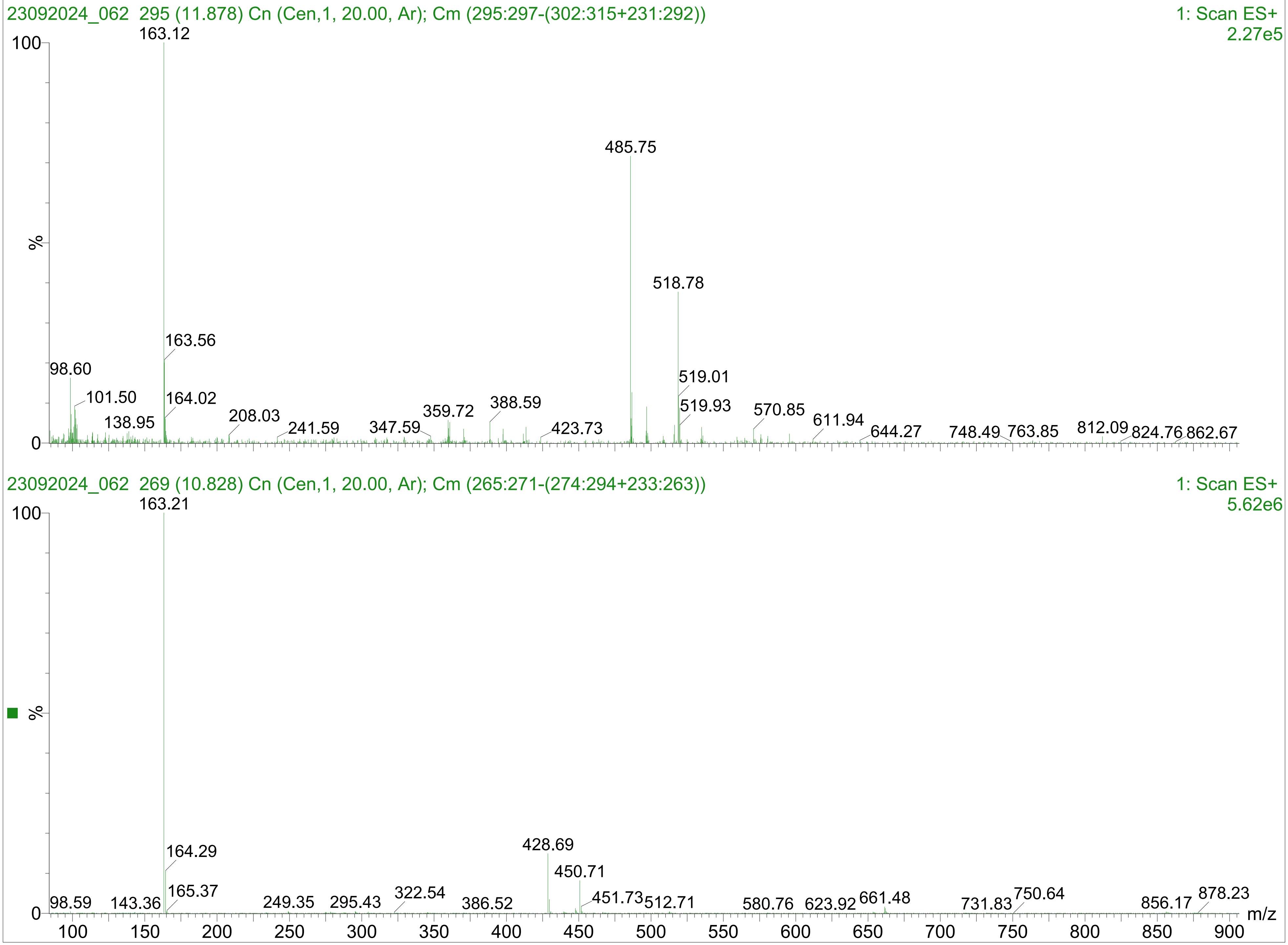

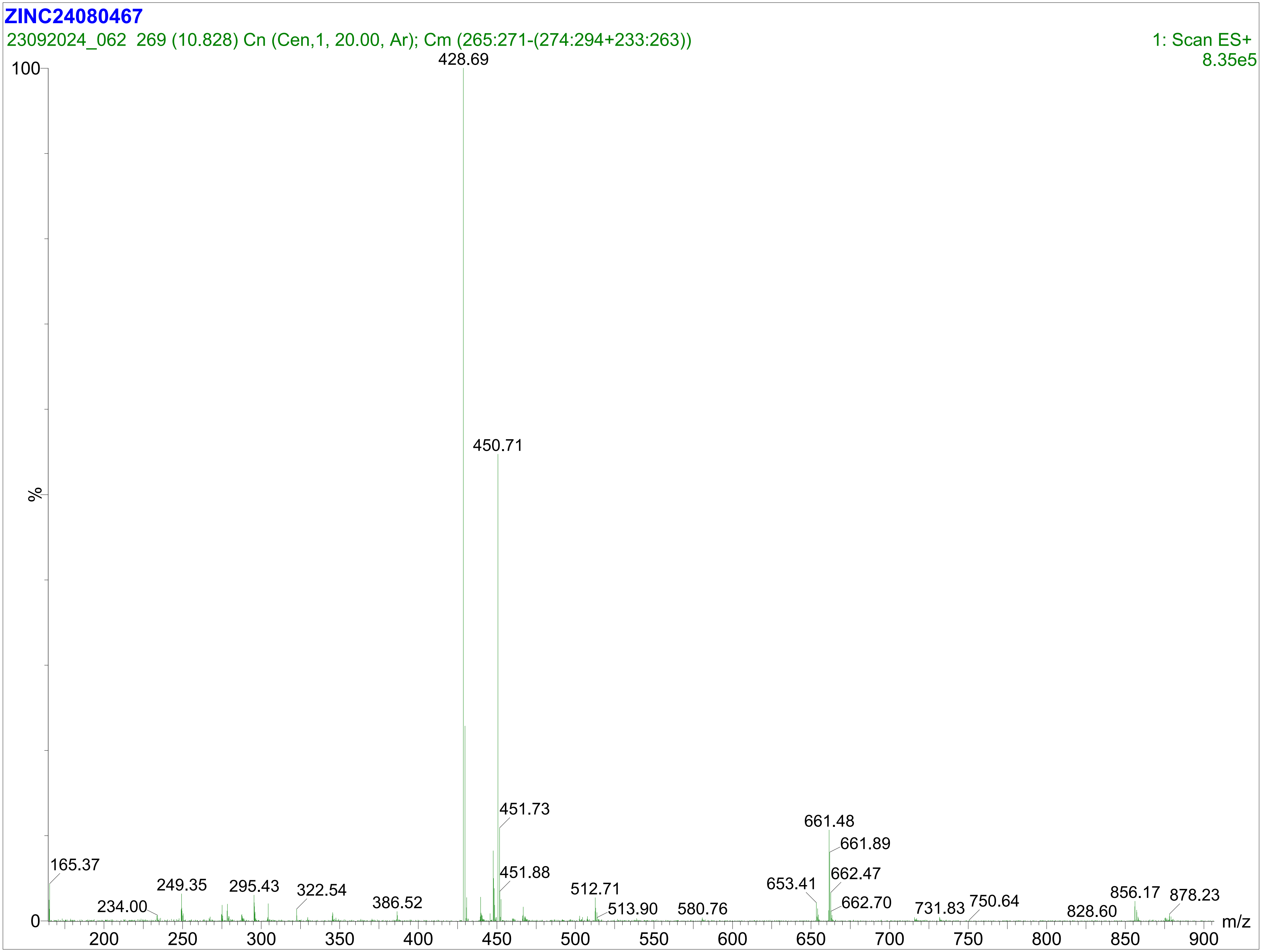
ZINC24080467
23092024_062 269 (10.828) Cn (Cen,1, 20.00, Ar); Cm (265:271-(274:294+233:263))
1: Scan ES+
8.35e5
100
%
0
428.69
450.71
451.73
451.88
165.37
234.00
249.35
295.43
322.54
386.52
512.71
513.90
580.76
653.41
661.48
661.89
662.47
662.70
731.83
750.64
828.60
856.17
878.23
200
250
300
350
400
450
500
550
600
650
700
750
800
850
900
m/z
